# Mapping 3D tissue orientation with tensor-augmented light-sheet microscopy

**DOI:** 10.64898/2026.05.05.722138

**Authors:** Mario Corral-Bolaños, Piyush Swami, Hanna Vila-Merkle, João Lima, Silja P. Vange, Christophe Destrieux, Cyril Poupon, Jessica Pingel, Sanne S. Kaalund, Lídia Bardia, Hans Martin Kjer, Marco Pizzolato, Jeppe Revall Frisvad, Julien Colombelli, Tim B. Dyrby

**Affiliations:** Department of Applied Mathematics and Computer Science, Technical University of Denmark, Kongens Lyngby, Denmark; Danish Research Centre for Magnetic Resonance, Department of Radiology and Nuclear Medicine, Copenhagen University Hospital Amager and Hvidovre, Hvidovre, Denmark; Imaging Brain and Neuropsychiatry iBraiN U1253, Université de Tours, INSERM, Tours, France; NeuroSpin, Paris-Saclay University, CNRS, CEA, Gif-sur-Yvette, France; Winsløw Unit of Anatomy, Institute of Molecular Medicine, University of Southern Denmark, Odense, Denmark; Centre for Neuroscience and Stereology, Bispebjerg University Hospital, Copenhagen, Denmark; Institute for Research in Biomedicine (IRB Barcelona), The Barcelona Institute for Science and Technology (BIST), Barcelona, Spain

**Author notes:** Corresponding author(s). E-mail(s): Tim B. Dyrby, Julien Colombelli.

## Abstract

Mapping the three-dimensional directional organization of biological tissues is essential for understanding their structure and function. However, existing methods cannot resolve micrometer-scale orientation in volumetric samples. We introduce tensor light-sheet scattering microscopy (tLSSM), a label-free method that reconstructs 3D fiber orientations at micrometer resolution in optically cleared tissues, enabling whole-organ imaging. We discovered that even in transparent samples, organized structures such as neural fibers scatter light in directional patterns, consistent with models of light scattering by cylinders. We compared tLSSM against diffusion MRI in the mouse brain, demonstrating strong agreement and orders of magnitude superior spatial resolution. Furthermore, we showcase tLSSM’s versatility across diverse contexts, including whole-brain label-free tracing, pathological demyelination lesions, heart tissue, peripheral nerves, and human white matter. By enabling whole-organ fiber orientation mapping compatible with standard light-sheet microscopes, tLSSM establishes a new standard for mesoscopic connectivity studies by mapping tissue architecture beyond the limits of traditional sectioning.

## 1. Introduction

Biological tissues are characterized by anisotropic microarchitectures, where the spatial orientation of cells and extracellular matrix components underpins physiological function. In the brain, neural circuits form a complex network with directional organization spanning scales from macroscopic inter-regional connectivity to microscopic axonal pathways^1^. This large-scale organization, termed macroconnectomics or projectomics, provides insights into how brain areas communicate and coordinate to support cognitive, behavioral and other functional processes^2^.

Mapping neural circuits remains a fundamental challenge. Early approaches, such as invasive tract-tracing, have enabled the construction of mesoscopic connectivity atlases from thousands of brains, such as the Allen Mouse Brain Connectivity Atlas^3^. However, these approaches based on one tracer injection per characterized tract are limited in scalability. Conversely, methods requiring no staining of tracers (i.e., label-free methods), like diffusion-weighted magnetic resonance imaging (dMRI), offer non-invasive, non-destructive tractography but suffer from a structural resolution limit^4^. Clinical dMRI voxels are typically millimeter-scale^5^, and even high-resolution ex-vivo setups remain orders of magnitude larger than individual axons^6–8^. This discrepancy leads to ill-posed orientation estimates and false-positive tract delineations^9,10^.

Efforts to bridge this gap have largely relied on tissue sectioning. Microscopic techniques such as polarized light imaging (PLI)^11,12^, serial optical coherence tomography (SOCT)^13,14^, and computational scattered light imaging (ComSLI)^15,16^ provide in-plane 2D orientation. Others, like polarization-sensitive OCT (PS-OCT)^17^, advancements on PLI^18,19^, or permittivity tensor imaging (PTI)^20^, offer 3D orientation information but remain limited to histological slices. Small-angle X-ray scattering tensor tomography (SAXS-TT)^21^ reveals 3D orientation of myelinated fibers in whole-mount (intact) specimens^22^, but high-resolution imaging is constrained by long acquisition times. Moreover, SAXS-TT and many of these techniques rely on myelin being intact, which may be compromised in pathological conditions.

Light-sheet fluorescence microscopy (LSFM) enables 3D volumetric imaging at micrometer resolution in intact optically cleared tissues. Clearing techniques homogenize refractive indices across tissue components (e.g., proteins, lipids, water), reducing light scattering and enabling deep imaging. LSFM has predominantly relied on fluorescent stains to visualize specific structures^23–25^. Recently, alternative light-sheet based studies have reported elastic scattering as a candidate for label-free imaging of small samples^26^, referring to the method as light-sheet tomography (LST), light-sheet scattering microscopy (LSSM), or scattered light-sheet microscopy (sLS). Elastic scattering originates from local refractive index variations, and it involves photons that are deflected without loss of energy. Elastic scattering provides information about the sample architecture across different scales: below the wavelength of incident light (Rayleigh scattering) and scales comparable to and larger than the wavelength (referred to as Mie scattering in biomedical optics)^27,28^.

Here, we present a 3D light-sheet scattering-based method for multiview mesoscopic imaging, enabling label-free, micrometer-resolution mapping of 3D fiber orientation in intact, optically cleared tissues. In this work, we refer to the acquisition of scattering data from multiple sample orientations as multiview light-sheet scattering microscopy (mvLSSM), and the subsequent directional reconstruction as tensor light-sheet scattering microscopy (tLSSM).

We demonstrate the application of tLSSM in the whole mouse brain, where we report that scattered light from neural fibers reveals an orientation-dependent signal, consistent with prior observations on collagen fibers^29^. To capture directional scattering, we developed an imaging workflow combining sample rotation, multiview acquisition, and 3D registration, allowing reconstruction of voxel-wise tissue orientation from multiple illumination angles. We demonstrate that, in cleared tissue, such a signal is (i) congruent with neural pathways, and (ii) capable of resolving the orientation of anisotropic tissue at sub-voxel resolution. We leverage the orientation dependency of the signal to propose a theoretical modeling framework that encodes orientation information in every voxel, enabling micrometer-scale 3D tractography. We verified that the estimated directions align with the dMRI-based results in the mouse brain. Finally, we explore tLSSM’s sensitivity to microstructure pathology in the brain and its applicability to other organs, such as the mouse heart, sciatic nerve and the human brain white matter.

## 2. Results

### 2.1. Scattered light-sheet microscopy reveals anisotropic signal in the brain

To characterize how the LSSM signal relates to the orientation of anisotropic tissue, we imaged iDISCO+-cleared mouse whole-brains at 4.5 µm isotropic voxel size using a dual-view, double-sided macroSPIM system (see Methods, Supplementary Methods 1 and Supplementary Figure 1 for instrument description).

Each imaging plane was acquired from four perspectives—LF, RF, LB, and RB—identified by light-sheet origin (left, L or right, R) and camera position (front, F or back, B). Unlike conventional LSFM, which assumes fluorescence is emitted isotropically^30^, LSSM reveals a structured, orientation-dependent signal absent in autofluorescence. This signal intensity is modulated by the light-sheet’s angle of origin, resembling neural pathways (Fig. 1b-e).

**Fig 1.**
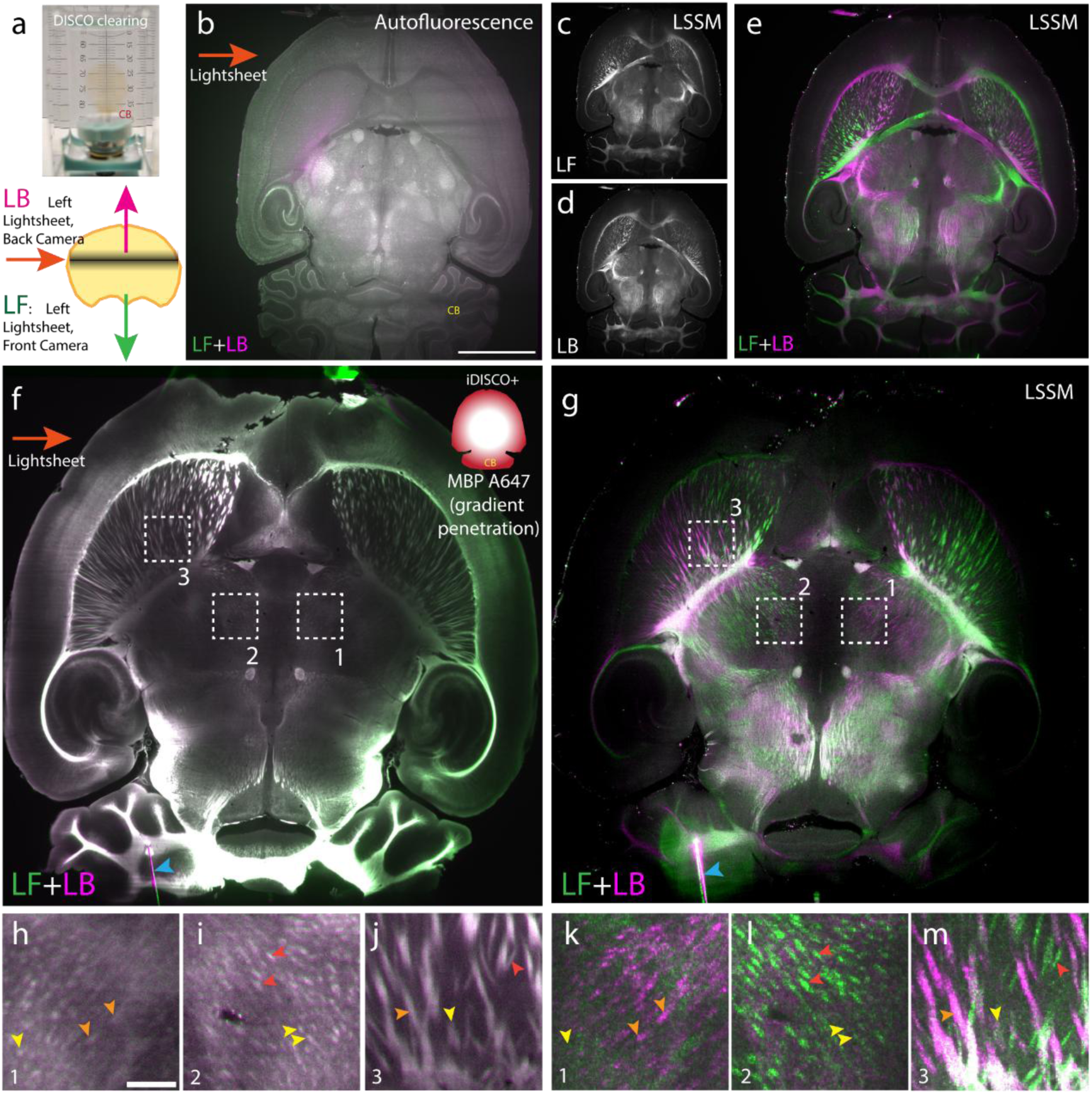
LSSM reveals axonal bundles in the mouse brain. **a.** (Top) Unlabeled mouse brain, cleared with iDISCO+, mounted vertically (CB: cerebellum down) on a silicon pad with pins (pin visible in **f-g**, blue arrowhead). (Bottom) Left light-sheet and detection side nomenclature, LB (back) and LF (front). **b**. Single slice (transverse, or axial) autofluorescence image showing the high similarity between both cameras (color merge LF green, LB magenta). **c-d**. scattered LSSM signal from the same light-sheet illumination from front (**c**) and back (**d**) cameras. **e**. Merged (**c**) (Green) and (**d**) (Magenta) showing the strong anti-symmetry in the collected signal depending on the detection side. **f**. Mouse brain (different sample from **b-e**) labelled with MBP-A647 (iDISCO whole mount labelling). Labelling efficiency decreases in the center (gradient due to penetration effects). Merged LF and LB showing similar fluorescence signals (green dominance can be interpreted as light-sheet illumination fading through the sample). **g**. Strong anti-symmetry of merged detected LSSM signals between cameras. **h-j**. fluorescence zoomed from dashed regions 1,2,3 in F. **k-m**. merged LSSM signals, same regions. Arrowheads point at signal foci: in red, MBP fibers with LSSM signal visible prominently on LF; in orange, signal visible on LB; in yellow, MBP fibers with no prominent LSSM associated signal. Scale: **b**, 2 mm; **h**, 0.2 mm.

To assess the anatomical relevance of the LSSM signal, we performed whole-mount immunofluorescence for myelin basic protein (MBP), a marker of myelinated axons forming the pathways. We observed that a single light-sheet perspective could not capture the full trajectory of the pathways (Fig. 1h-m and Extended Data Fig. 1). Instead, complementary patterns emerged across opposing perspectives (Fig. 1c-e), and similarities appeared between crossed pairs with opposite cameras and light-sheets (Extended Data Fig. 1). These patterns suggest the signal is anisotropic and modulated by the relative amount of collected scattered light, depending on the combined orientations of the light sheet and the detection axis. LSSM signals co-localized with MBP fluorescence in regions such as the corpus callosum and striatal fibers in the putamen (Fig. 1f,h,i,j vs. Fig. 1g,k,l,m), supporting the interpretation that LSSM highlights neural pathways, albeit in a directionally dependent manner (Fig. 1h-m).

To further understand the structural length scale of LSSM signal origin, we tested the signal dependence on linear polarization orientation (see Supplementary Note 1 and Supplementary Fig. 1f-p). Switching to out-of-plane (p-pol, perpendicular to the image plane) from in-plane (s-pol, parallel) polarization decreased background but largely suppressed fiber-associated signals (Supplementary Fig. 1o). A polarization-independent signal persisted at the brain surface, likely from large particles. The polarization-dependent signal from fibers is consistent with Rayleigh scattering^31^, suggesting it could be induced by cellular structures at scales below the light wavelength (here <638 nm). Whereas the brain surface LSSM signal is more consistent with a Mie scattering behavior, a regime occurring at, or above, the light wavelength scale by larger debris, such as dust particles that are more polarization-invariant.

Importantly, we observe that the scattering signal intensity depended strongly on tissue processing. Fiber-associated scattering was prominent in paraformaldehyde-fixed brains but markedly reduced with glyoxal fixation (Extended Data Fig. 1y–ag). In contrast, similar signals were observed across solvent-based (BABB/iDISCO+) and water-based (CUBIC) clearing methods in our sciatic nerve analysis (Supplementary Note 2 and Supplementary Fig. 2,3). This indicates that while the scattering signal may be preserved across clearing protocols, fixative chemistry plays a critical role in scattering visibility.

### 2.2 Characterization of the directional scattering signatures of the 3D neural pathways

To characterize scattering signals across different positions in a cleared mouse brain, we acquired an image series of volumetric data across a full 360° in-plane rotation range in nominal 20° steps (i.e., a mvLSSM dataset). The resulting 3D volumes were registered to a common space using an in-house algorithm (Supplementary Methods 4). The intensity changes with rotation steps were particularly evident in white matter structures, such as the corpus callosum and internal capsule (Fig. 2a,b, see also Supplementary Videos 1,2,3).

**Fig 2.**
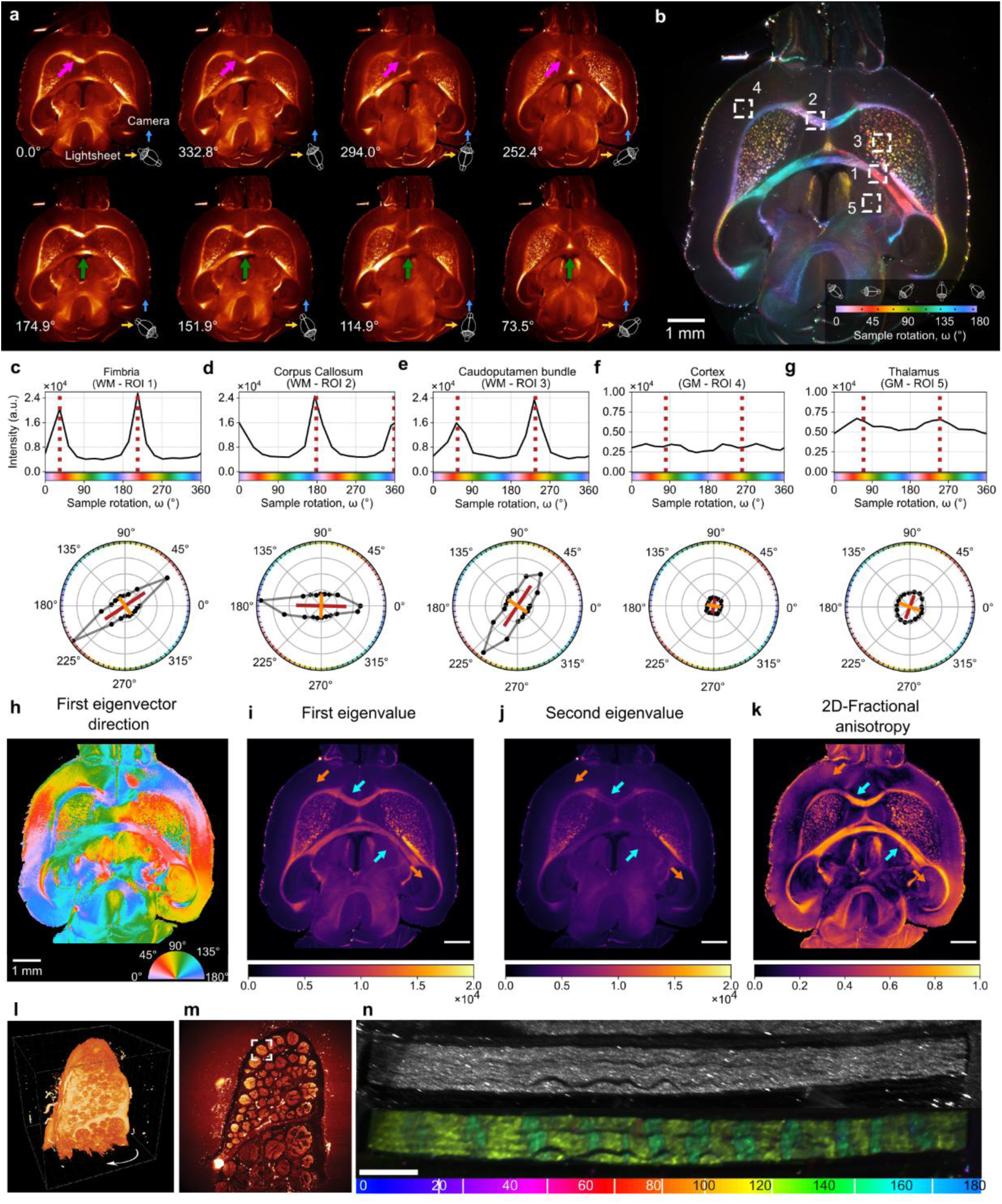
Rotational characterization of the scattering signal. **a.** Representative axial slice showing rotation-dependent contrast across white matter structures (magenta and green arrows). The estimated rotation angle applied during registration is indicated in the bottom-left corner of each panel. **b.** Pseudo-colored composite image representing the LSSM contrast variations across sample rotation angles in the [0, 180)° range. The pseudocolor of each volume depends on its applied rotation, as indicated in the bottom-left legend. The black markers indicate the specific color of each volume. The center of the white squares indicates selected regions of interest for subsequent analysis. **c-g.** Top row: mean scattering profiles from five 6×6×1 voxel regions of interest in white matter tracts (c, d, e) and gray matter (e, f), with estimated peak positions marked by dashed brown lines. The x-axis indicates the right-hand rotation applied to the sample. Bottom row: corresponding profiles projected into 2D polar space, showing first (brown) and second (orange) principal directions. **h-k.** Maps of the (**h**) first eigenvector orientation, (i,j) first and second eigenvalue, and (k) 2D fractional anisotropy characterizing scattering profiles across the entire slice. Cyan arrows indicate white matter structures; orange arrows indicate gray matter regions. **l-n**. Horse sciatic nerve axon bundle. (l) 3D rendering of the horse sciatic nerve section. (m). Scattering channel from a single view, the white square indicates the cross section of the bundle shown in (n). (n) top: single-view scattered signal. (b) pseudo-colored composite image representing the mvLSSM contrast variations across sample rotation positions. Scale: b, h-k) 1 mm; n) 0.5 mm

Voxel-wise scattering signal profiles were extracted from the image series (Fig. 2c-g). In white matter regions containing densely packed axons, such as the corpus callosum and fimbria, the profiles showed two peaks approximately 180° apart (179.98 ± 1.57°), reflecting the symmetry of fiber scattering (Fig. 2c-e). In contrast, gray matter regions such as the cortex and thalamus displayed broader, less distinct peaks, likely reflecting the absence of a predominant direction, yielding a more isotropic signal. Still, the 180° periodicity could be appreciated (Fig. 2f,g).

To quantify peak position and its prominence, we used the principal component analysis (PCA) algorithm. PCA was applied voxel-wise on a 3D space (Fig. 2c-e, bottom row), where each data point represents the applied rotation to the sample and the measured intensity in each voxel. The first eigenvector represents the applied rotation to measure a peak (Fig. 2h). Notably, scattering peak positions acquired from opposing light-sheet-camera pairs (LF/RB vs. RF/LB) were consistently offset by 90° (Extended Data Fig. 2a,b). The PCA first eigenvalue (Fig. 2i), representing peak amplitude, was highest in white matter regions (cyan arrows) independently of their orientation, highlighting densely packed axon bundles such as the corpus callosum and caudoputamen striale fibers, respectively. The second eigenvalue (Fig. 2j), sensitive to baseline intensity and peak width, showed patterns similar to those of the first eigenvalue. The degree of signal anisotropy (i.e., the fractional anisotropy, FA), derived from the normalized ratio of the first and second eigenvalues, was high in white matter (cyan arrows) and lower in gray matter (orange arrows) dominated by somas, suggesting tissue type sensitivity in light-scattering (Fig. 2k).

We confirmed the symmetry of the scattering profiles by comparing the PCA-features from fitting only half-range ([0, 180)°) and the full-range (360°) data (Extended Data Fig. 2c-g). We obtained consistent results in all the features with the two approaches, with some small deviations on the first eigenvector in some gray matter areas, probably due to the lack of prominent peaks.

Finally, for obtaining clear insights on the scattering signal relationship with the orientation of the fibers, we imaged a horse sciatic nerve, which contains densely packed, parallel axonal bundles (or fascicles) that serves as a reference sample with easy-to-retrieve macroscopic straight orientations (Supplementary Note 2 and Supplementary Fig. 2-4). Beyond the macroscopic straight direction of the axonal bundles, mvLSSM revealed mild local undulations along the axonal trajectories. These local variations, known as Bands of Fontana, can be appreciated in the scattered signal (Fig. 2n). Notably, mvLSSM highlighted striped bands at different rotation positions during acquisition. These varying scattering-contrast bands, corresponding to the rising and falling edges of the axonal undulations, support the high sensitivity of mvLSSM to mesoscopic trajectory variations (Fig. 2m).

### 2.3 Modeling the 3D fiber orientation using infinite-cylinder scattering theory

To reconstruct the 3D fiber orientations from a mvLSSM dataset, we modeled light scattering from fibers as infinite cylinders, parameterized by two angles (Fig. 3a): the in-plane azimuthal orientation (β_f_) and the out-of-plane tilt (α_f_). Theoretical predictions based on Maxwell’s equations indicate that light is scattered along a cone centered on the fiber axis, with the cone’s aperture determined by the angle between the incident light propagation direction and the fiber orientation (Extended Data Fig. 3a-f, Supplementary Methods 3, Supplementary Video 4)^32^.

**Fig. 3.**
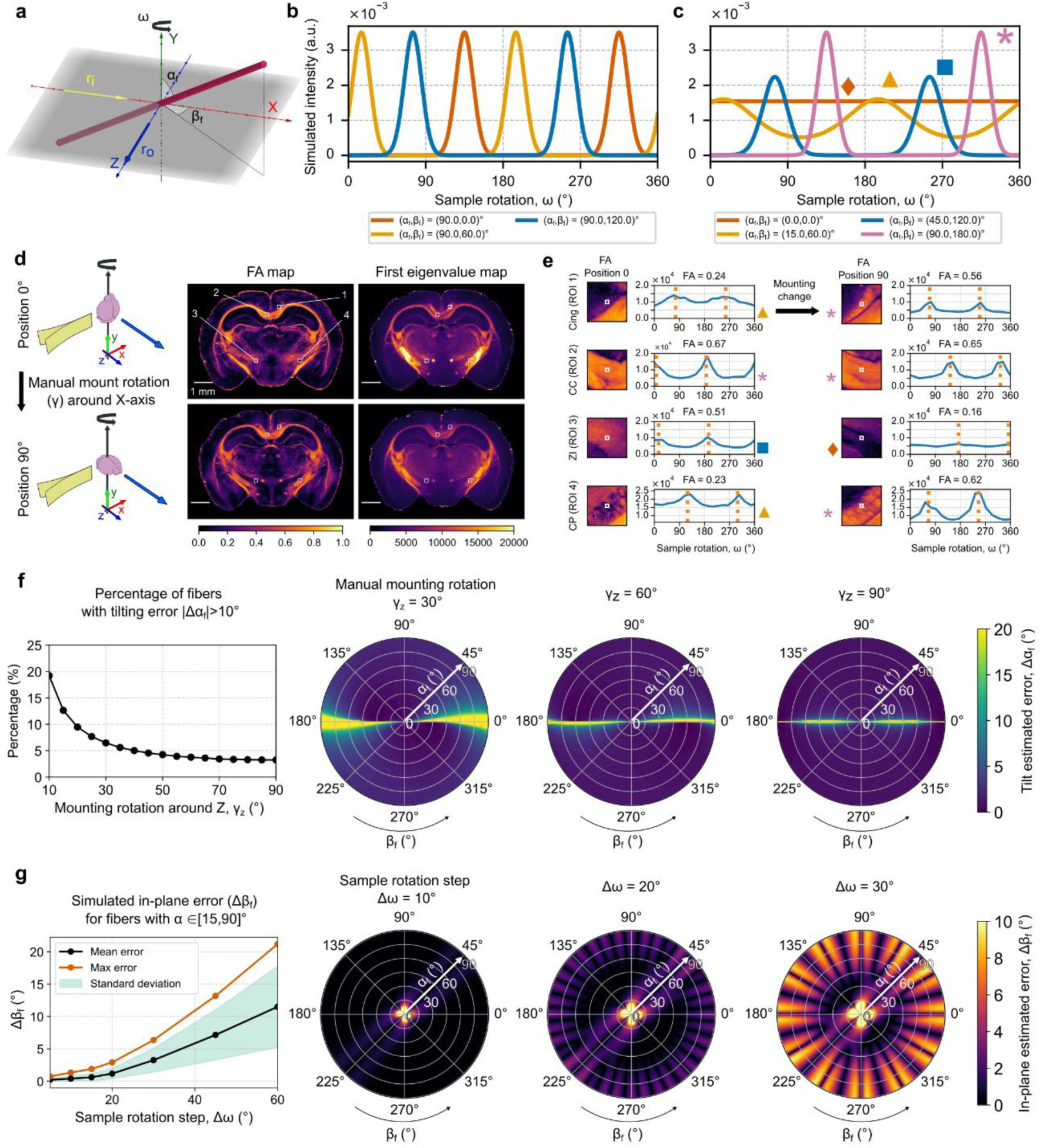
Theoretical and experimental evaluation of the infinite-cylinder model and acquisition considerations. **a.** Schematic of a 3D fiber modeled as an infinite cylinder, defined by tilt (αf) and azimuthal (βf) angles, ω, ri and ro denote the sample rotation direction, light-sheet trajectory and camera position, respectively. **b,c.** Simulated scattering profiles in the macroSPIM system, legends indicate the initial fiber orientation: **b**, in-plane fibers (αf=0°) with different initial azimuth βf=0,60,120°. **c.** fibers with varying tilting, αf. The peak position represents the necessary rotation for a fiber to be placed at β=45°. **c.** Experimental FA and first eigenvalue maps obtained from mvLSSM with the sample at two different mounting orientations (top position 0, bottom position 90). The left schematic shows the approximate position of the brain mounting. **d.** Comparison of the scattering profiles across different ROIs (8×8×1 voxels) along with a zoom-in of the FA map in the two mounting positions. Regions: Cingulum (Cing), Corpus Callosum (CC), Zona Incerta (ZI), Caudoputamen fibers (CP). The icon between scattering profiles links simulated amplitude modulation (b, right) to experimental data in each mounting. **e.** Simulated effect of the manual rotation between the mvLSSM datasets around the Z axis on the out-of-plane estimation. The polar plot is interpreted as the top-view of a hemisphere representing all the possible fiber directions. The radius indicates tilt angle and the polar angle indicates the azimuth. Left: percentage of orientations with tilt error >10° versus rotation angle. Right: tilt error across orientations for selected mounting-rotation angles. **f.** Simulated effect of the angular sampling step-size on the in-plane orientation (βf). Left: mean β error across sampling steps. Right: β error for selected sampling resolutions.

We first simulated scattering profiles for the macroSPIM configuration, where the rotation axis (Y-axis) is perpendicular to both the illumination (X-axis) and detection axes (Z-axis). When simulating a sample rotation, we are effectively changing the in-plane, β, orientation of the fiber. The first insight is that a peak only appears when the in-plane orientation component is β=45° (Fig. 3b). On the other hand, the peak amplitude is expected to vary with tilting (α_f_ > 0°), such that out-of-plane fibers show lower peak amplitudes than in-plane fibers (Fig. 3c). When the fiber is aligned with the rotation axis (α_f_=0°), the scattering intensity profile is constant. All these insights are aligned with the experimental observations in the sciatic horse nerve study (Supplementary Note 2 and Supplementary Fig. 2).

While our study is focused on the macroSPIM system, we also assess other light-sheet hardware configurations. We simulated the scattering profiles for the ultramicroscope configuration, where the detection is placed on the top of the sample. The effective difference with respect to the macroSPIM (that features a side detection), is that the rotation axis is the same as the detection axis (i.e., the Z-axis). Importantly, this causes the relationship between fiber orientation and scattering profile features to be different (Extended Data Fig. 4). We further discuss the ultramicroscope system in Supplementary Note 3.

We then verified the macroSPIM simulations with experimental data, where the peak position of the signal profile robustly represents the in-plane orientation (β_f_). However, the peak amplitudes (reflected in the first PCA eigenvalue, Fig. 2i) did not vary substantially with out-of-plane fiber orientation, indicating challenges in correlating amplitude with fiber tilt. Nonetheless, we still observed some changes in the peak prominence (reflected in the 2D-FA map) with respect to its baseline in specific areas, as predicted by simulations. To confirm this observation, we manually rotated the brain by 90° (in practice, approximately 75°), and compared the 2D-FA maps for each sample mounting position (Fig. 3d). Differences between the two FA maps were only evident in areas where the fibers were approximately aligned with the rotation axis (α_f_=0) in one of the two mountings. For example, in the posterior cingulum (Fig. 3d, region of interest 1), the 2D-FA was noticeably larger in one of the mountings than the other.

To better understand the limitations of the amplitude modulation, we simulated potential confounding factors, such as variations in fiber diameter or orientation distributions (Extended Data Fig. 3g-j). The simulations confirmed that such features could influence the amplitude modulation, complicating its use as a reliable proxy for fiber orientation. Other factors, such as fiber density^33^, or practical differences with the theoretical model, such as light attenuation, the angular dispersion on the incident illumination or the numerical aperture in detection, may further spread the scattering peaks and reduce orientation specificity.

### 2.4 Two-planes acquisition resolves the complete 3D fiber orientation

To robustly recover full 3D fiber orientation, we combined two mvLSSM datasets, with the second obtained after manual sample rotation (∼75°) about an axis in the XZ plane (Fig. 3d). This yields two independent in-plane orientation measurements, β and β′, from which the out-of-plane tilt (α_f_) can be estimated via triangulation avoiding the use of the amplitude modulation in a single rotation-plane dataset.

Simulations confirmed that any non-zero manual rotation between mvLSSM datasets enables the estimation of α_f_ (Fig. 3f). However, a small subset of fiber tilting orientations remains ambiguous depending on the second mounting orientation (Methods; Supplementary Video 5). For example, for Z-axis rotation between mountings, fibers near the XY plane (β_f_ ≈ 0°/180°) are affected by tilting uncertainty. The subset of affected fibers decreased the closer the manual rotation was to 90°. Still, we observed that <5% of orientations would be affected for inter-acquisition rotation angles >40°, well below the >75° used here (Fig. 3f).

We also evaluated the acquisition rotation step (Δω). Simulations using observed white-matter peak widths (∼40°) showed that Δω = 20° balances accuracy and acquisition time, yielding mean β_f_ errors <5% (Fig. 3g). Higher uncertainty occurs for fibers aligned with the rotation axis (flat profiles) or with large tilt (α_f_ ≈ 90°) due to narrower peaks, respectively. Experiments with Δω = 10° and 20° produced consistent results across white-matter ROIs (mean difference 2.00 ± 2.06°; Extended Data Fig. 5).

### 2.5 Comparative evaluation of tLSSM with diffusion MRI for fiber orientation mapping in the mouse brain

We compare the 3D fiber orientation from the single-fiber tensor fit of the two-plane mvLSSM datasets (i.e., tLSSM) and well-established diffusion tensor MRI on the same brain. The volume ratio between tLSSM and the dMRI data acquired using a preclinical 7T MRI system is approximately 28^3^-fold higher: isotropic 4.5 μm vs. 125 μm resolution. For tLSSM, data from all camera/light-sheet pairs were combined to improve angular resolution, and to promote consistent results in gray matter, where the lack of prominent peaks increases sensitivity to light-sheet attenuation caused by optical path differences. Nevertheless, single-pair reconstructions yielded consistent results in white matter (Extended Data Fig. 6). Tissue shrinkage introduced by the clearing protocol (estimated ∼30% per axis) was compensated using tensor-based affine and non-linear registration to align mvLSSM with the dMRI volume. Finally, results were rigidly rotated to align with the Allen Mouse Brain Connectivity Atlas^3^.

Overall, the RGB color-encoding (see Methods) of fiber orientation showed strong correspondence between tLSSM and dMRI (Supplementary Video 6). A small fraction of the voxels (≈0.02%) were affected by tilt ambiguity due to the triangulation method and were solved through neighbor information. Major axon-dense white matter structures, such as corpus callosum, cingulum bundle, fimbria, and optic tract, are clearly resolved by both methods, as their cross-sectional area exceeds a dMRI voxel (Fig. 4a-d, cyan arrows). Minor artifacts were observed in regions with crossing fibers, such as the projections of the caudoputamen fibers into the corpus callosum, arising from the model assumption of a single fiber orientation per voxel when using the PCA-based fitting (Fig. 4i).

**Fig. 4.**
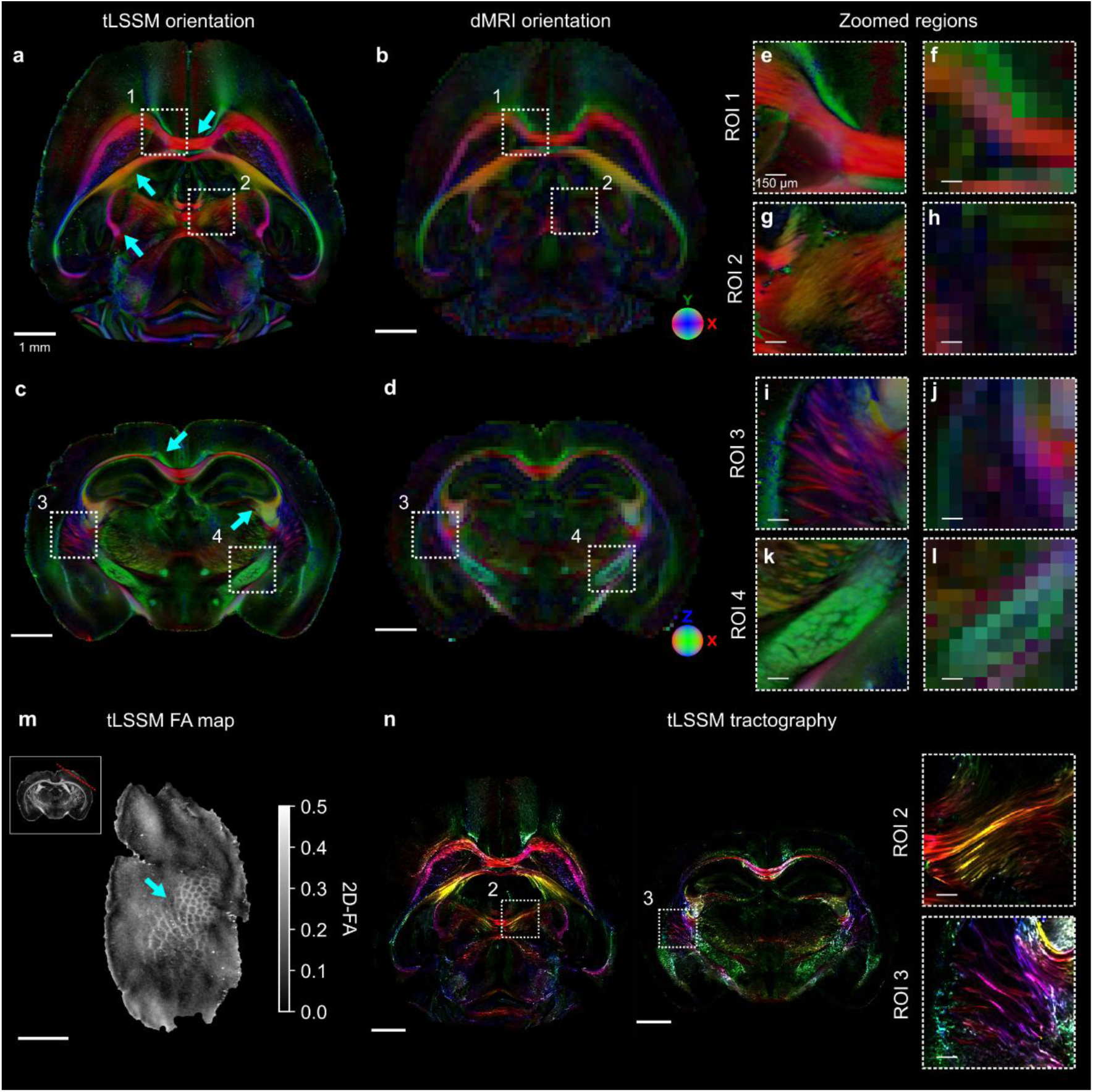
Comparison of fiber orientation and structural resolution between tLSSM and dMRI in the mouse brain. **a–d**. RGBA color-coded directionality maps from (**a,c**) tLSSM (left) and (**b,d**) dMRI (right), showing major axon-dense structures (cyan arrows). **e–l**. Zoomed views of selected regions: corpus callosum (**e,f**), thalamic projections (**g,h**), striatal fascicles in the putamen (**i,j**), and internal capsule (**k,l**). For each pair, tLSSM is shown on the left and dMRI on the right. **m.** tLSSM-fractional anisotropy map of the barrel field (cyan arrow). On the top left, location of the barrel field in the mouse brain (red line). **n**. Track-density imaging reconstruction based on tLSSM-derived fiber directions, and zoom-in of the reconstructed tracts in the same region as in **g** and **i**. Here, higher saturation represents a larger number of tracks going through each voxel, and the color-hue represents the orientation as in a-l. **Scale**: a) 1 mm, e) 0.150 mm.

tLSSM resolved axon bundles and individual somas beyond the structural resolution limit of dMRI. In the corpus callosum, tLSSM reveals stripes of axon bundles likely navigating around glial clusters, visible as low-intensity bands^34^, not resolved by dMRI due to its low resolution (Fig. 4e,f). In the internal capsule, typically mapped as a unified white matter tract in dMRI and atlases, tLSSM revealed smaller striatal axonal bundles interwoven with deep gray matter structures (Fig. 4k,l). Similar but thicker striatal bundles are seen in the putamen, partially captured by dMRI (Fig. 4i,j). tLSSM also resolved thin axon bundles in the lateral posterior nucleus of the thalamus and brain stem-region, undetectable by dMRI (Fig. 4g,h).

In the cortex, both methods show lower regional anisotropy probably reflecting the high cell density and lack of a predominant fiber direction as seen in Fig. 2k. Notably, tLSSM captures representative cortex regions non-distinguishable in dMRI, such as the barrel field (Fig. 4m), appearing as low-anisotropy domains separated by the higher-anisotropy septa, reflecting the underlying somatotopic organization.

To demonstrate label-free virtual path tracing for mesoscale structural connectomics, we performed whole-brain streamline-based tractography on mouse tLSSM data in Figure 4a,c. Streamlines seeded in random voxels were propagated by following local tLSSM fiber orientations. In the resulting directionally encoded track-density map (see Methods, Fig. 4n; Supplementary Video 7), color hue determines the average streamline orientation, while color saturation reflects the density of reconstructed tracks. The visualization reveals white-matter pathways with RGB-coded directionality consistent with tLSSM orientation maps (Fig. 4a,c), resolving axonal fascicles in the corpus callosum and fine separate bundles of striatal fiber projections in the putamen and brainstem (Fig. 4n, zoom-in), in agreement with local tLSSM features (Fig. 4g,i).

### 2.6 Applications of LSSM in pathology, human brain and heart tissue

We induced focal axonal demyelination in the mouse corpus callosum by injecting 1% LPC in one hemisphere, with a sham injection contralaterally. tLSSM revealed reduced anisotropy within the lesion, reflected by lower peak amplitude in fiber profiles (Fig. 5a-c), consistent with decreased MBP fluorescence labeling in the same brain confirming myelin loss. Hematoxylin and eosin (H&E) staining confirmed an LPC-induced immune response inflammation with increased cell density (Fig. 5d)^35^, likely contributing to the reduced signal scattering in lesion area. Notably, axonal orientation was preserved as expected^1^, and tLSSM still detects fiber orientations within the lesion, although the FA values were lowered.

**Fig. 5.**
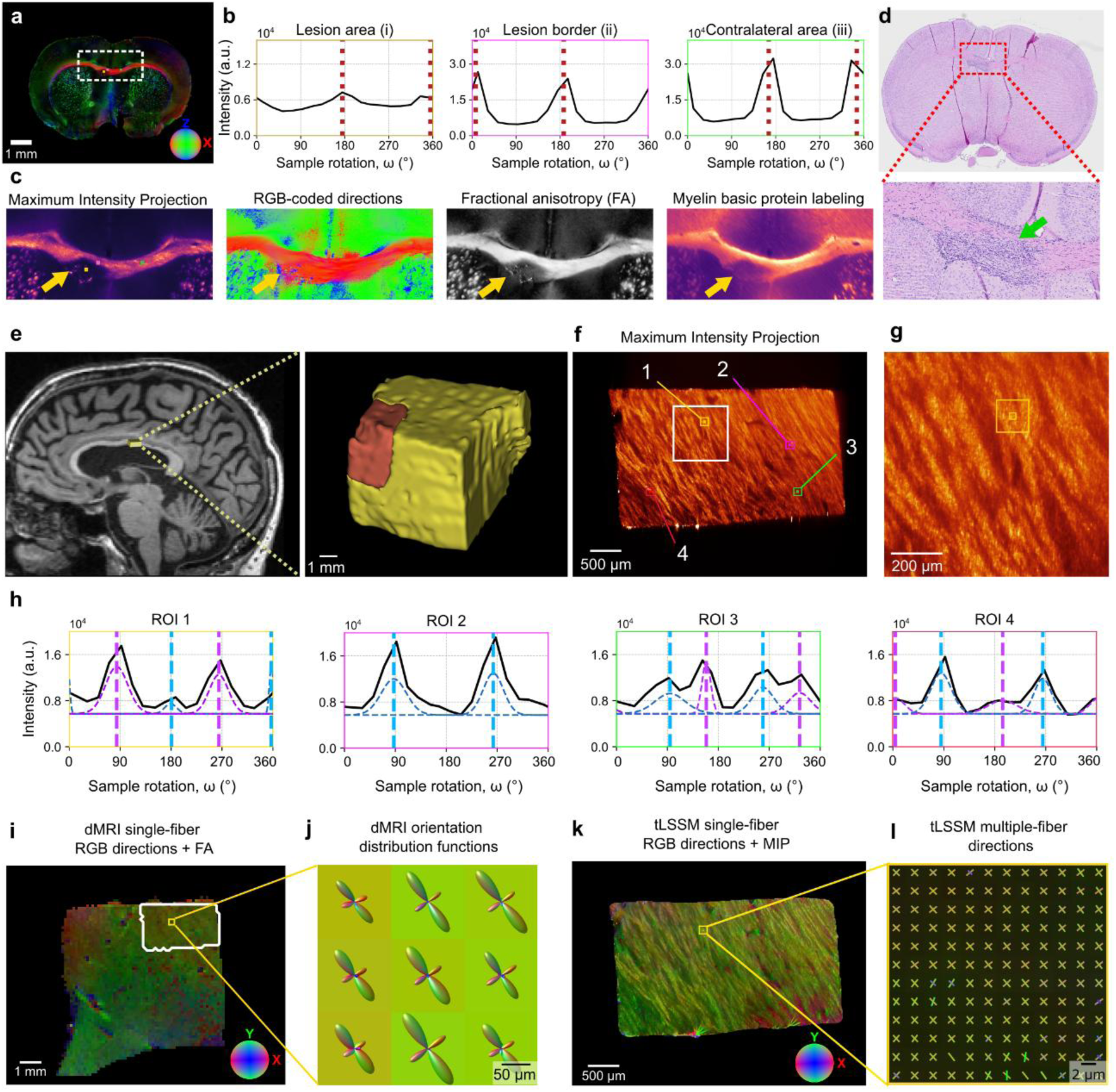
Applications of tLSSM in pathological mouse and healthy human brain. **a.** tLSSM reconstruction of fiber orientations in a mouse brain with focal demyelination induced by 1% LPC in the corpus callosum (orange arrow), with a sham injection in the contralateral hemisphere. **b**. Mean scattering profiles from 3×3 voxel regions across lesion and control areas. **c**. Zoomed tLSSM-derived maps showing fiber directionality and anisotropy, alongside MBP fluorescence. **d.** H&E staining on a LPC-induced lesion (green arrow) on a second mouse brain. **e.** In vivo MRI of the subject with the approximated area of the imaged human corpus callosum piece and its 3D rendering. The entire piece was imaged with dMRI, while the orange region was imaged with mvLSSM. **f.** Maximum intensity projection of the mvLSSM acquisition. **g.** Zoomed region of the composite image showing white matter bundles. **h.** Averaged scattering profiles over the 10x10x1 voxel ROIs depicted in (f) revealing different configurations. **i,k.** RGB-coded single-direction fiber orientations derived from dMRI (i) and tLSSM (k). **j,l.** Crossing fibers orientation revealed by dMRI (j) and tLSSM (l) in approximately the same region.

In a cleared human corpus callosum sample (Fig. 5e), mvLSSM revealed stacked axonal layers with distinct orientations (Fig. 5f,g and Extended Data Fig. 7), including crossing fibers characterized with a von Mises mixture fiber model-based algorithm (Fig. 5h). Crossing fibers were robustly detected at 2.46 μm isotropic resolution, consistent with diffusion MRI at 125 μm (Fig. 5i-l)^36^ and supported by X-ray synchrotron data from monkey brain at 75 nm resolution^1^. Across regions of interest, scattering profiles revealed crossing fibers with varying peak amplitude, likely reflecting differences in the density of fibers within each bundle (Fig. 5g).

Finally, mvLSSM applied to a cleared mouse heart also revealed directional scattering, reflecting cardiomyocyte network complexity (Extended Data Fig. 8a-c), where profiles of both single and crossing fibers were detected (Extended Data Fig. 8d,e)^37^, demonstrating the mvLSSM method’s applicability beyond neural tissue.

## 3. Discussion

We demonstrated that the intrinsic scattering properties of optically cleared tissues can be decoded to map 3D fiber orientation at micrometer scale in whole, intact organs. By developing tensor light-sheet scattering microscopy (tLSSM), we addressed the resolution limits inherent to dMRI^4^ while avoiding the scalability constraints of tracer-based or sectioning methods^2^. We showed the versatility of tLSSM across diverse biological architectures, including the intact mouse brain and heart, horse sciatic nerve, and human white matter. As a result, we have established a scalable and non-destructive framework for mapping anisotropic tissue architecture, enabling the virtual tracing of whole-brain projections at the micron axon-fascicle level and offering sensitive markers for microstructural pathology.

To assess the robustness of the method, we compared tLSSM with dMRI fiber orientation maps on the same mouse brain (∼0.5 cm³). We found that overall orientation patterns are stable across modalities and robust to averaging artifacts when a voxel contains multiple tissue types (i.e., partial-volume effects)^1^. While we observed strong hemispheric symmetry and closely matched diffusion tensor MRI–derived and tLSSM orientation maps (Fig. 4a-d), we achieved an approximately 21,400-fold (∼28³) increase in volumetric resolution, enabling us to reveal complex architectural details that are blurred at dMRI scales^4^. For example, we clearly resolved striatal axon fascicles in the caudate and putamen and detected fascicular projections into brainstem pathways that are not visible with MRI (Fig. 4a-l). Similarly, we captured anatomically relevant variations in gray matter anisotropy, such as the mouse barrel cortex (Fig. 4m). This fidelity enabled us to perform label-free, streamline-based tractography in individual brains that closely resembled the tracer-based maps of the Allen Mouse Brain Atlas^3^. The achievable image resolution of tLSSM is primarily limited by the optical field of view (1.44× magnification) required for whole-mouse brain coverage. Further improvements toward submicron resolution are possible at higher magnification using 3D image stitching. Higher resolution can also be obtained with smaller fields of view, as we demonstrated here in the sciatic nerve and human corpus callosum. Notably, even at isotropic 2.46 µm resolution in the human corpus callosum, crossing fibers were inferred, a structural feature consistent with dMRI^36^ and X-ray synchrotron imaging findings by Kjer et al. in the monkey corpus callosum^1^.

We attribute the origin of the tLSSM elastic scattering contrast to the anisotropic arrangement of subcellular structures. The marked signal reduction upon altering illumination polarization is consistent with a Rayleigh-scattering-dominated regime^31^ (Supplementary Note 1), implying sensitivity to structures significantly smaller than the illumination wavelength (<638 nm), such as neurofilaments (∼10 nm)^38^, microtubules (∼25 nm)^39^, or lipid bilayers (∼5 nm)^40^. These elements maintain cellular architecture and, when averaged within voxels, would reflect tissue-level anisotropy. This interpretation is supported by the persistence of directional signals in the sciatic nerve: since the expected mean axon diameter (7 µm)^41–43^ exceeds the imaging voxel size (0.7 × 1.5 × 1.5 µm^3^), the observed intra-voxel anisotropy must originate from the alignment of subcellular elements rather than the axon boundaries (Supplementary Figure 3i-w). Prior histological work has also identified distinct subcellular structures as major scatterers, including organelles rich in membranes^44^ and periodic protein assemblies (sarcomeres in muscle tissue)^45,46^. Unlike fluorescence, which targets specific proteins, elastic scattering integrates the cumulative arrangement of these subcellular elements, serving as a proxy for tissue-level anisotropy. This mechanism is consistent with our observations in other anisotropic tissues, such as the sarcomere alignment in cardiomyocytes, suggesting broader applicability to structurally ordered tissues.

Beyond orientation mapping, we leveraged the sensitivity of subcellular structures to detect microstructural alterations in focal demyelinating lesions. Voxel-level anisotropy reflects an ensemble average of the scattering profiles from a mixture of microstructural compartments such as cells and axons. Consequently, cellular-dominated regions like the cerebral cortex exhibit a more isotropic scattering profile. This aligns with observations in diffusion MRI, despite the fundamental difference in probing mechanisms (light scattering vs. water diffusion). This averaging effect also elucidates signal changes in pathology; in focal demyelinating lesions of the mouse brain, we observed reduced anisotropic scattering (Fig. 5a-d). This is possibly driven not only by myelin loss but also by an inflammatory response characterized by increased density of infiltrating glial cells^35^, which introduces isotropic scattering signal—a mechanism supported by recent synchrotron and MRI imaging studies at 0.5 and 200 µm resolution, respectively^1,47^.

We identified the sample preparation protocol as a critical factor for preservation of the scattering signal. We observed that while the scattering signal is robust across solvent- (iDISCO+, BABB) and water-based (CUBIC) clearing protocols, and compatible with immunofluorescence labeling (MBP staining), it is sensitive to fixation chemistry. The scattering signal was preserved in PFA-based perfusion fixed tissues but reduced in glyoxal-fixed samples. While the underlying mechanism requires further study, we hypothesize that the distinct protein cross-linking network formed by PFA may be essential for preserving the refractive index fluctuations that generate the scattering contrast^48^.

Furthermore, we showed the generalizability of the theoretical framework of tLSSM to other light-sheet microscopy configurations. However, the reconstruction requires accounting for the specific geometry of the system; for example, the use of different orthogonal systems (e.g., macroSPIM vs. Ultramicroscope) is conceptually equivalent to altering the sample’s rotation axis (Fig. 3a,d vs. Extended Data Fig. 4a,b). Consequently, the specific features of the scattering profile—and how they encode fiber orientation—vary, needing system-specific modeling (Supplementary Note 3). Currently, robust 3D reconstruction relies on a two-plane acquisition strategy to resolve the intrinsic confounding factors between fiber tilting and signal amplitude. Brain fibers are not straight but form orientation and radii distributions^1,49^, which confound the intensity variations that could be used to estimate out-of-plane angles. While we mitigated this through manual rotation of the sample and triangulation, this increases acquisition time and data volume. Future iterations incorporating alternative optical designs could achieve single-mounting 3D reconstruction, simplifying the workflow.

Ultimately, we envision tLSSM as a powerful synergistic tool with spatial biology. As a non-destructive, label-free modality compatible with fluorescence, tLSSM could be integrated with whole-mount immunostaining or emerging whole-brain spatial transcriptomics studies^50^. This would allow researchers to overlay molecular maps (e.g., cell types, gene expression) onto a precise 3D structural scaffold. Moreover, our findings in cardiac and pathological tissue indicate this versatility extends beyond neuroscience. We anticipate that tLSSM will enable the development of multimodal atlases across diverse organ systems, providing a versatile tool to unravel the complex relationship between microscopic tissue anisotropy and molecular function in health and disease.

## 4. Methods

### 4.1 Tissue preparation

Animal models and tissue preparation procedures are detailed on Supplementary Methods 5. Here, a brief description is provided.

#### Mice

A total of 7 mice were used in this study. From those, 6 were used for mvLSSM imaging and 1 for Hematoxylin and Eosin (H&E) staining. To induce focal demyelination in the corpus callosum of adult wild-type mice (n=2 BomTac:NMRI females; Taconic Biosciences, Denmark), an intracranial injection of 1 μL of 1% lysophosphatidylcholine (LPC) in sterile saline was administered. Seven days later, lesion size was verified using in-vivo MRI. Mice were euthanized with an overdose of pentobarbital and transcardially perfused with 4% paraformaldehyde (PFA) or glyoxal in heparinized saline. Brains of LPC-injected (n = 1) and healthy (n = 5) mice and one heart (n = 1) were then extracted and stored at 4°C. All these samples underwent mvLSSM imaging, and one healthy brain underwent ex-vivo MRI imaging previous to clearing.

#### Horse sciatic nerve

After euthanasia, the horse sciatic nerve was dissected from the left hindlimb, sectioned proximally to its bifurcation into the N. tibialis and N. peroneus communis, embedded in 4% PFA, and stored at 4°C for tissue clearing.

#### Human

Data from a 94-year-old male participant without any known neurological medical history, who was included in the FIBRATLAS project, were used for this study. The patient died at the age of 98 years. Five hours after death, both common carotid arteries were injected with one liter of 4% buffered formalin. The brain was extracted 17 hours after death, it was hung by the basilar artery into 4% buffered formalin and stored at 4°C to slow down autolysis^51^. After fixation, a median sagittal cut was made to separate both hemispheres. The brain was stored at 4°C for 3.5 years. Finally, the small piece of the corpus callosum used in this study was cut and stored in PBS at 4°C during 5 days for posterior MRI imaging, clearing and mvLSSM.

### 4.2 Diffusion magnetic resonance imaging

After fixation, the brain was placed in PBS for rehydration at least one week prior to MRI to restore the T_2_ signal^6,51^. Diffusion MRI data were acquired ex vivo from a single mouse brain using a 7T Bruker Biospec MRI scanner at the Danish Research Centre Magnetic Resonance (DRCMR), Copenhagen University Hospital Amager and Hvidovre, Denmark equipped with a dual-channel cryogenic coil. The acquisition protocol employed a single-shell pulsed-gradient-spin-echo (PGSE) sequence with a b-value of 8000 s/mm^2^ (δ = 6 ms, Δ = 12 ms), comprising 64 non-collinear diffusion encoding directions and five b=0 s/mm^2^ volumes. Images were acquired with an isotropic voxel size of 0.125 mm, echo time of 25 ms, and repetition time of 5.7 s. To reduce susceptibility-induced distortions, the brain was immersed in a perfluorocarbon solution (Fluorinert®) during scanning^6^. Single fiber orientations were estimated using the diffusion tensor imaging model^52^, and fiber orientation distribution function (fODF) were reconstructed with the Constrained Spherical Deconvolution (CSD)^53^.

For the human subject, an in vivo MRI scan was performed on a 3T Siemens Prisma scanner, including a T1-weighted scan (GR\IR, TE: 2.98 ms, TR: 2300 ms, voxel size 1×1×1 mm).

### 4.3 Optical clearing and immunolabeling

Following fixation (and MRI scanning), non-labeled tissue samples (mice brains, mouse heart and human corpus callosum) were processed using the iDISCO+ clearing protocol^54^. Samples were dehydrated at room temperature through a graded methanol (MeOH) series (20%, 40%, 60%, 80%, and 100%), with each step lasting ∼1 hour. Delipidation was performed in a 66% dichloromethane (DCM) 33% methanol solution for ∼3 hours at room temperature, followed by two washes in 100% DCM. Tissues were then incubated in dibenzyl ether (DBE), which also served as the final imaging medium.

Samples labeled for myelin basic protein (MBP) underwent a pretreatment step to enhance antibody penetration, followed by immunolabeling with a fluorophore-conjugated mouse anti-MBP primary antibody (see Supplementary Methods 5 for details). Subsequently, the same iDISCO+ clearing protocol was applied.

For the horse sciatic nerve, the same sample was used for comparative clearing with solvent-based MeOH:BABB and water-based CUBIC clearing (see Supplementary Methods 5).

### 4.4 Multi-view light-sheet scattering microscopy imaging

Light sheet imaging was performed at the Barcelona Mesoscopic Imaging Node of Eurobioimaging-ERIC, in the Advanced Digital Microscopy facility of IRB Barcelona. The employed imaging system was a custom-built selective plane illumination microscopy based on double-sided macroscope detection (macroSPIM) and double-sided illumination, previously applied to whole organ imaging in fat pad^55^, liver^56^, and bladder tissues^57^. We upgraded it to include light-sheet polarization control to modulate and maximize scattered signal, axially swept light-sheet illumination microscopy (ASLM) to maximize axial resolution, and temporal pivot scanning to mitigate coherent laser speckle (see Supplementary Methods 1 and Supplementary Fig. 1 for full instrument description and performance). Scattered signal was recorded with a 638 nm laser. mvLSSM acquisitions of mouse brains, mouse heart and human sample were imaged at 1.44× magnification yielding a voxel size of 4.514 μm^3^, with ASLM (9 or 11 bands), while sciatic nerve was imaged at 1.92× (voxel 3.385 μm^3^), without ASLM (static light-sheet); see supplementary Note 1 for datasets details. mvLSSM data (mouse brain, sciatic nerve, heart, human brain) was acquired from 18 angular views (step approximately 20°) with a custom magnetic rotation base, and when needed the sample was manually rotated by 90° to acquire a second 18-views dataset.

To evaluate the scattering signal patterns through a different rotation-axis, we utilized the Ultramicroscope II (UMII, Miltenyi/LaVision BioTec GmbH, Germany), which had top-mounted detection and required no customization or modification (see Supplementary Note 3 for details). Acquisitions were performed using left-sided illumination and a 620 nm laser. Brain samples were imaged at 1.26× magnification (0.63× optical zoom). Whole-brain 3D volume acquisition was conducted across 16 views via planar rotation using a custom-built sample holder.

### 4.5 Registration of light-sheet scattering volumes

To enable voxel-wise modeling of the scattering profile, a multi-resolution image registration pipeline was implemented using the Elastix software package^58^ (see Supplementary Note 4 for detailed description). Volumetric images were downsampled by a factor of four in each dimension to improve computational efficiency, and gamma correction (γ=0.5) was applied to reduce the influence of directional dependent high-intensity scattering regions.

Affine transformations were estimated by maximizing mutual information (32 histogram bins) between fixed and moving images. Registration was performed at three resolutions using a smoothing pyramid (downsampling factors: 4,2,1). All volumes were ultimately registered to a reference volume acquired at “0°” (the first chosen sample orientation) mounting with the back camera and left light-sheet.

The registration proceeded in two stages: (1) intra-series alignment of each volume I_n_ to I_n-1_, followed by composition of transforms to align all volumes to I_0_; and (2) inter-series alignment to the reference volume. Final refinement was performed at the original voxel-size using 2000 iterations without pyramid smoothing.

### 4.6 Registration of diffusion MRI to the LSSM space

To align mouse brain diffusion MRI data to the LSSM space, fractional anisotropy (FA)^59^ maps were generated for both modalities to facilitate registration. The LSSM data were downsampled to 90 μm isotropic voxel-size. An initial affine registration using normalized correlation and a three-level smoothing pyramid (factors: 4,2,1) was applied. This was followed by a non-linear B-spline registration using normalized correlation (weight 1) and a bending energy penalty (weight 0.5), with control point spacing of 250 μm and up to 500 iterations. For the human white matter sample, a manual rigid registration was performed to align both acquisitions. The resulting transformations were applied to the preprocessed dMRI data.

### 4.7 Light scattering by infinite cylinders modeling

Neural fibers were modeled as infinitely long circular cylinders to simulate light scattering under light-sheet illumination. For this model, the interaction and scattering of a plane electromagnetic wave with wavelength shorter than the cylinder length is well described^32,60^ (see Supplementary Methods 3 for a detailed description). When light is incident on an infinite-long cylinder, the scattered light propagates along the surface of a cone, with its apex at the point of incidence. The cone’s half-angle is given by π/2 - φ, which matches the angle between the incident light direction and the fiber orientation.

The solution to the scattering problem requires identifying the scattered field in any point of space. To this end, we used the open-source MATLAB package MatScat^61^. The total intensity in each point of the scattered cone depends on the elevation angle, φ; and the scattering angle, θ. The angle φ is related to the aperture of the cone of scattered light, and θ identifies an angle around the cylinder’s cone, i.e., it identifies a particular propagation direction within the cone.

Unless otherwise stated, the scattering intensity was simulated with the parameters^62^: incident wavelength λ = 572 nm, cylinder radius ρ_0_ = 8 nm, cylinder refractive index N = 1.407, medium refractive index N_m_ = 1.562, and relative permeability between the outer and inner medium μ_i_/μ = 1. To match acquisition settings, macroSPIM simulations (Y-axis rotation) simulated incident s-polarized light, and ultramicroscope simulations (Z-axis rotation) simulated incident unpolarized light.

#### Scattering profile simulation

In the microscope coordinate system (XYZ), the model predicts a measurable signal intensity when the cone of scattered light (r_s_) is tangential to the detector normal **r**_o_, i.e., **r**_s_ = **r**_o_ (see Extended Data Fig. 3a-d). Thus, given the definition of the elevation angle **r**_f_ ⋅ **r**_i_ = cos(π/2-φ) = sin(φ) and given that the detector normal r_o_ is perpendicular to the light-sheet (**r**_i_ ⋅ **r**_o_ = 0). We can obtain θ using the angle between two planes:

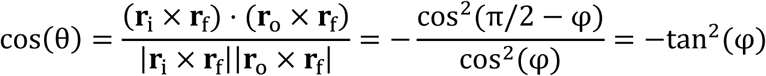

Once φ and θ are known, the expected scattering amplitude is given by the scattering function.

The variations in axonal trajectories along their length give rise to a fiber orientation distribution with a predominant direction^49^. Thus, the scattering profiles were modeled as weighted integrals over the fiber orientations. Fiber orientations, r_f_, were parameterized using spherical coordinates defined by two angles: α_f_, the tilt relative to the rotation axis, and β_f_, the azimuthal angle in the perpendicular plane. Each voxel’s predominant fiber direction was used to compute the expected signal S(α_f_, β_f_) by integrating the scattering function S(α,β) over a distribution of orientations. The probability distribution 𝒫(α,β|α_f_,β_f_) was modeled as a normal distribution weighting the acute angle between the main fiber direction, α_f_,β_f_, and the direction, α,β, 𝒩(0,σ^2^), where the standard deviation, σ, was set to 10° unless otherwise stated.

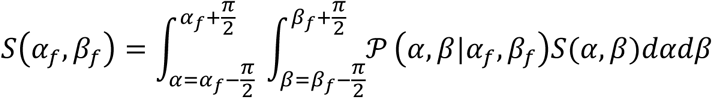

Simulations were performed using system-specific configurations. For consistency, the light-sheet propagation was set in the X-axis, **r**_i_ = (1, 0, 0); and the detection in the Z-axis, **r**_o_ = (0, 0, 1). Then, for macroSPIM, the rotation axis was set to (0, 1, 0); while for the ultramicroscope system, the rotation axis was set to (0, 0, 1). The resulting scattering amplitudes were computed using the scattering intensity S(φ, θ).

### 4.8 Principal component analysis for peak detection

By leveraging the periodicity of the peaks in the scattering profile, the orientation of the peaks can be accurately determined. First, we extract the applied rotation from the registration pipeline. We then assign the coordinates (1, 0, 0) to a reference volume and determine the relative positions of the other volumes based on the applied rotation.

Next, we map the scattering profiles to the 3D space using spherical coordinates. For each value of the scattering profile, we assign the polar and azimuthal coordinates corresponding to their volume, while the radius represents the voxel intensity. Then, we compute the two first principal components voxel-wise. The direction of the first eigenvector represents the necessary rotation to be applied to the sample to measure the maximum intensity. It allows us to estimate peak positions with a smaller angular resolution than the actual angular step.

Additionally, the PCA approach enables us to quantify of the peaks prominence relative to a baseline using the 2D-Fractional anisotropy^59^:

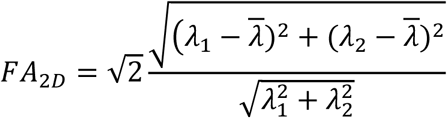

The values of FA range from 0, when λ_1_=λ_2_, case of a homogeneous distribution of the intensities; to 1, when λ_1_>>λ_2_, indicating high peak prominence. We evaluated the accuracy of the PCA by simulating the scattering profiles with varying angular sampling resolutions. This was done for all possible fibers with values α ∈ [0, 90]° and β ∈ [0, 360)°, using 1° spacing between values. We tested the sampling resolutions 5°, 10°, 15°, 20°, 30°, 45°, and 60°.

We assessed the experimental angular periodicity of the scattering profiles using a circular autocorrelation approach applied voxel-wise via the fast Fourier transform. The profiles were resampled to 9° increments, and periodicity was defined as the lag of the highest normalized autocorrelation peak exceeding 0.10. Experimental periodicity was reported as the mean ± standard deviation within a white matter region of interest on a single axial slice, restricted to voxels with fractional anisotropy larger than 0.4.

### 4.9 The inverse problem: reconstructing the 3D fiber orientation

The inverse problem for reconstructing 3D fiber orientations includes two steps. Firstly, we should find the in-plane orientation of the fibers in two planes, namely β and β’. Afterwards, the 3D fiber orientation can be solved as a computer vision triangulation problem. The approach for the first step may vary when using hardware configurations (i.e., macroSPIM or ultramicroscope), while the second is applicable to any case. See Supplementary Note 3 for notes on the ultramicroscope system.

#### In-plane fiber orientation in macroSPIM

The azimuthal orientation of fibers within the XZ plane was inferred by analyzing the modulation of scattering profiles as the sample was rotated around the Y-axis. According to the theoretical model, αf influences the amplitude and peak width, βf relates to the lateral shift of the scattering profile. Due to limited sensitivity to tilt in the experimental data, sensitivity to noise and potential confounding factors, we focused on extracting βf from the peak position. Since we know that peaks only happen when the in-plane fiber position is placed at 45°, and the effect of rotating the sample is to change the in-plane fiber position, we can recover the original fiber orientation is:

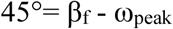

where ω_peak_ represents the right-hand rotation around +Y applied to the sample to measure a peak.

#### 3D fiber orientation as a triangulation problem

To resolve the out-of-plane orientation, we leveraged the flexibility of LSSM to acquire data from multiple sample mountings. By manually rotating the sample and repeating the acquisition, we obtained scattering profiles from two non-parallel planes.

After 3D registration, the fiber orientation rf was reconstructed via triangulation from its projections **r**_fa_ and **r**_fb_ on planes A and B, as the intersection of the two planes with normal (**r**_fa_ × **n**_A_) and (**r**_fb_ × **n**_B_), respectively.

The 3D triangulation works if the planes (**r**_fa_ × **n**_A_) and (**r**_fb_ × **n**_B_) are not parallel. To evaluate the prevalence of this case, we simulated fibers with α ∈ [0°, 180°) and β ∈ [0°, 180°) at 0.25° resolution and applied manual rotations around the z-axis. The tilting resolution, Δα, was defined as the fraction of fibers that, while initially sharing the same azimuthal angle β, exhibited a transformed azimuth β’ after manual rotation such that the angular difference |β’ - β| remained within a specified tolerance Δβ. More formally,

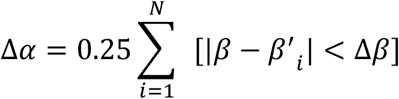

where [] is the Iverson bracket, and i indicates all the fibers with the same β.

### 4.10 Tensor-LSSM streamline based tractography

Tractography was performed using the fiber assignment by continuous tracking (FACT) algorithm^63^. A total of 30 million streamlines were generated using deterministic tracking, with default step size, maximum angle deviation 60°, and minimum tLSSM-Fractional anisotropy 0.1. Tracking was seeded uniformly across the brain. Track-density imaging (TDI) was performed by mapping the number of generated streamlines passing through each voxel onto the original imaging resolution^64^. All processing and visualization steps were conducted within the open-source software MRtrix3.

### 4.11 Directionally encoded color FA and TDI maps

For both tensor-based and streamline-based visualizations, the estimated local fiber orientations were mapped to the red-green-blue (RGB) color space. The absolute directional components of the orientation vectors were assigned as follows: red for the x-axis (left-right), green for the y-axis (anterior-posterior), and blue for the z-axis (superior-inferior). In the case of FA maps, the raw RGB values representing vector direction at each voxel were multiplied by the scalar FA value, rendering isotropic regions dark and anisotropic regions bright^65^. In the case of TDI, the assigned RGB color was determined by calculating the average direction of all intersecting streamlines passing through the voxel. To reflect tract density, the color saturation was modulated by the number of streamlines passing through the voxel^64^.

## Data availability

A sample dataset including one-shell DWI, registered mvLSSM and fitted tLSSM macroSPIM data from the same mouse brain will be publicly available upon publication at BioImageArchive (https://www.ebi.ac.uk/biostudies/studies/S-BIAD2486).

## Code availability

The code for simulations and data fitting will be publicly available upon publication at https://github.com/MaP-science/FibreSpy.

## Ethical declarations

All experimental procedures on mice were approved by the Animal Experiments Inspectorate of Denmark (Ethical protocol: 2023-15-0201-01494) and conducted in accordance with national and international regulations for the care and use of laboratory animals.

The sciatic nerve tissue samples were taken from a horse that was donated to the Department of Veterinary and Animal Sciences, University of Copenhagen by private owners after euthanasia for reasons unrelated to this study. The horse was euthanized by a captive bolt pistol followed by exsanguination. Informed consent was obtained from all owners, and the procedures are approved (2018-15-0201-01462) by the Animal Ethics Institutional Board, Department of Veterinary and Animal Sciences, University of Copenhagen. None of the procedures are considered animal experiments under Danish law, and national permits were therefore not required.

The human tissue samples were taken from a participant registered to Tours University’s body donation program before participating in the Fibratlas Project. Informed and written consent was obtained from him, and he consented to the terms under which his body would be used for academic or research purposes. The Fibratlas study adhered to the Ethical Principles for Medical Research Involving Human Subjects laid down in the Declaration of Helsinki and was approved by the Tours Ethics Committee (Comité de Protection des Personnes, 2015-R8) and by the ANSM (Agence Nationale de Sûreté du Médicament et des Produits de Santé, EudraCT/ID RCB: 2015A00363-46). The authors state that every effort was made to follow all local and international ethical guidelines and laws that pertain to the use of human cadaveric donors in anatomical research.

## Acknowledgements

The project was carried out with funding provided by the Lundbeck Foundation (grant number: R370-2021-1010), and research grant (17492) from VELUX FONDEN. The project has received funding from the European Research Council (ERC) under the European Union’s Horizon Europe research and innovation programme (grant agreement No. 101044180, ‘CoM-BraiN’) (Principal Investigator: T.B.D). This work received funding from Danmarks Frie Forskningsfond (Denmark Independent Research Fund) with grant ID 3105-00129B (Principal Investigator: M.P.) and grant ID 10.46540/4286-00333B. The human specimen collection was supported by the French National Research Agency (Fibratlas ANR-14CE17-0015, Summit ANR-21-CE45-0022-03, and France 2030 PEPR-SN BrainDeepPhenotyping ANR-22-PESN-0011). The authors thank the participant who donated his body to science so that anatomical research could be performed. Results from such research can potentially increase mankind’s overall knowledge, which can then improve patient care. Therefore, this donor and his family deserve our gratitude. We are thankful for the laboratory assistance of Vibeke Høgsberg.

## Author contributions

T.B.D. conceived the study and coordinated the experiments. J.C. built/adapted the macroSPIM instrument for LSSM and acquired the light-sheet data. J.C. analyzed sciatic nerve data. M.C.B acquired and analyzed the ex-vivo magnetic resonance imaging data. M.C.B derived the orientation reconstruction model with input from J.R.F. M.C.B., P.S., H.M.K and T.B.D. developed the registration pipeline. M.C.B. wrote the simulation, orientation reconstruction and analysis software. S.P.V. extended the implementation to crossing fibers. H.V.M., J.L., and T.B.D. prepared tissue samples and performed the tissue clearing of mice brain and heart, and human white matter. H.V.M and J.L. performed the LPC surgeries. H.V.M. performed the MBP labeling. S.K and J.L. performed the H&E histology. J.P. prepared the sciatic nerve tissue. L.B. performed the tissue clearing of the sciatic nerve. C.D and C.P. acquired the in-vivo human data and prepared the human brain tissue. M.C.B. analyzed the data with input from M.P. and T.B.D. M.C.B and J.C. prepared the figures. M.C.B. wrote the original draft with inputs from J.C. and T.B.D. Research was supervised by T.B.D. The text was reviewed and edited with input from all the coauthors.

## Additional information

Supplementary information is available for this paper. Correspondence and requests for materials should be addressed to Tim B. Dyrby or Julien Colombelli.

---

**Extended Data Fig. 1.**
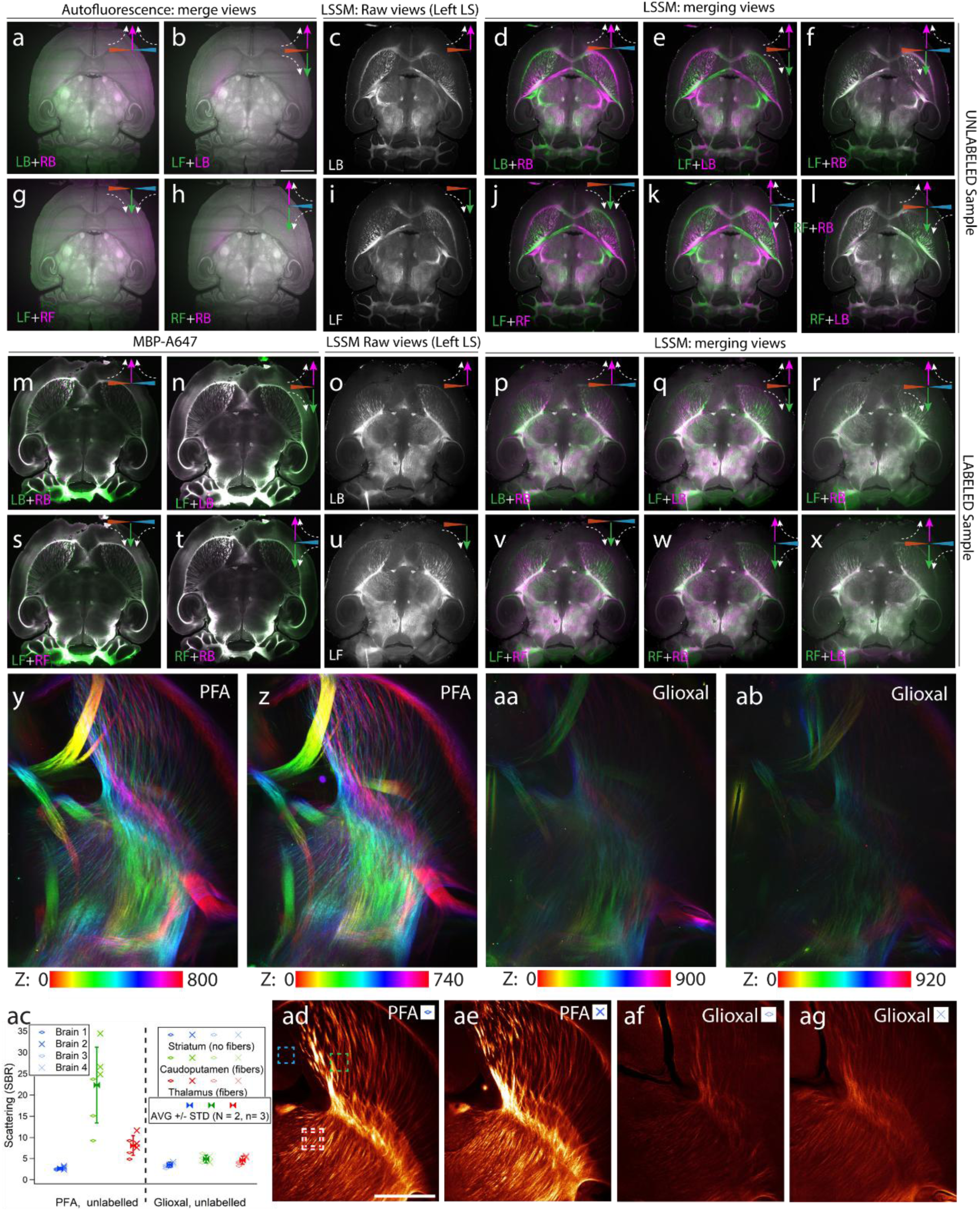
Anisotropic scattered signal emanating from brain fibers and its redundancy (extended data of Figure 1) and effect of brain fixation on scattered signal. **a-l.** Unlabeled mouse brain. **a,b,g,h.** Autofluorescence signal, merges of dual sided detection and dual sided illumination views showing similar images with minor differences due to light-sheet fading and/or opposite light-sheets misalignment. **c,i.** LSSM, Left Front and Left Right respectively. LSSM, merges from opposite LS illumination (**d,j**) or opposite detection side (**e,k**) show antisymmetric/complementary signals. Merging crossed view LF+RB (**f**) or LB+RF (**l**) shows similar signal distribution, affected again by light-sheet fading and/or non-perfect light-sheets alignment. **m-x.** In a MBP647 labelled mouse brain, merged views show the same similarities in fluorescence (**m,n,s,t**), complementarity of LSSM signal in opposite views (**o,p,u,v**) and similar LSSM signal in crossed views (**r,x**). **y-ag.** LSSM signal depending on fixation. **y,ab.** depth-color coded volume in a region including striatum, caudoputamen and thalamus (axial depth reported below images, in μm). PFA fixation in 2 brains (**y,z**) vs. Glioxal fixation (**aa, ab**). Whole mouse brains imaged at 1.44× magnification in **a-x**, whole brains at 4.8× in **y,z,aa,ab** and corresponding zoomed views **ad-ag** (4 different animals). **ac.** Signal-to-Background intensities are measured in n=3 small squares of interest (dashed, only 1 represented per region) in the regions striatum, caudoputamen and thalamus for each brain (N=2 for PFA, N=2 for Glioxal). For each brain region, averages (over N=2 brains) of scattered LSSM signal are reported with standard deviation (vertical bars).

**Extended Data Fig. 2.**
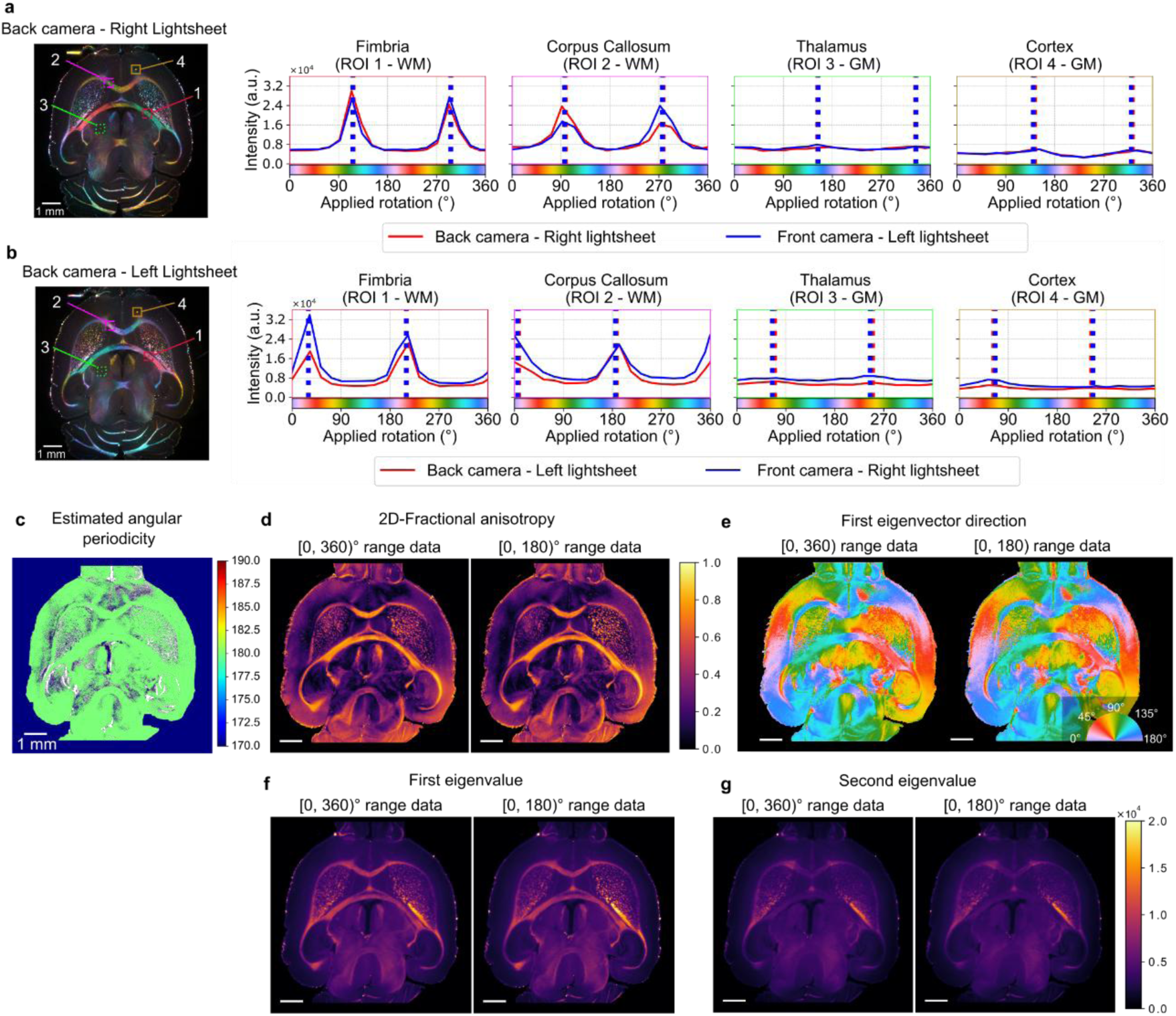
Redundancy analysis of scattering profiles from mvLSSM. **a-b.** Pairwise comparison of detector and light-sheet combinations. On the left, a pseudo-colored composite mvLSSM dataset acquired over a 180° range. On the right, scattering profiles from four regions of interest (ROIs), with estimated peak positions. Equivalent combinations yield consistent peak estimates, with a systematic 90° shift observed between **a** and **b**. Differences between the first and second peaks within the same profile suggest light-sheet degradation due to optical pathway differences. **c.** Estimated angular periodicity of the scattering profile through circular autocorrelation. **d-f.** Maps of 2D fractional anisotropy (**d**), eigenvector orientation (**e**), and first (**f**) and second (**g**) eigenvalues computed using all volumes (left) and only those within the [0, 180)° range (right). Results are consistent across both approaches, revealing the 180° periodicity. Minor deviations, particularly in cortical regions with low prominent peaks (e.g., orange arrow in **e**), could reflect reduced signal quality or light-sheet degradation.

**Extended Data Fig. 3.**
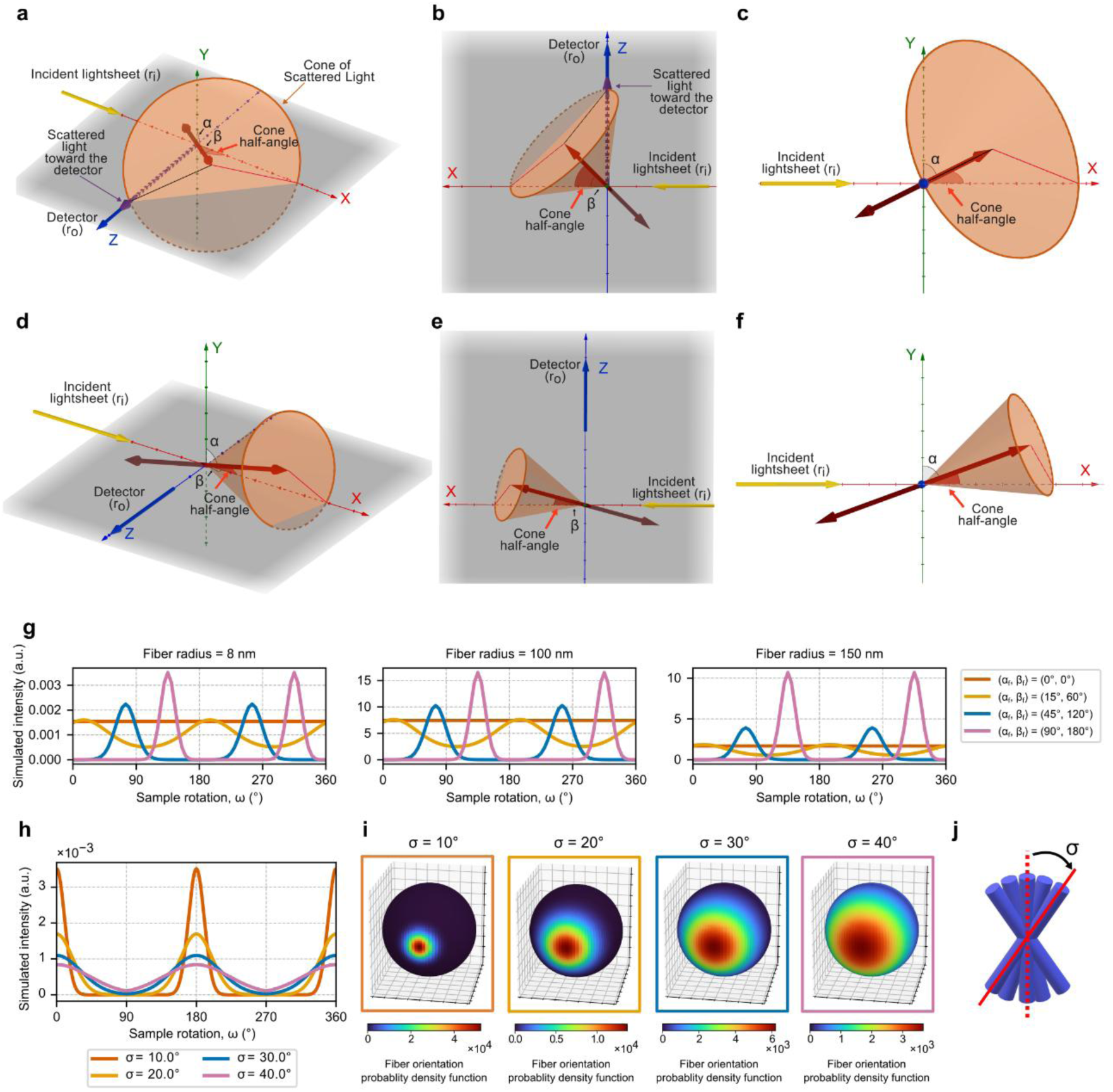
Relationship of the fiber position with the scattering peaks and confounding factors on the peak-intensity. **a-f.** Scattering cone generated by a fiber, rf, and main elements in the microscope setup. The angle between the light-sheet propagation and the fiber is the cone half-angle. **a-c.** The fiber orientation (α,β) = (70.5, 45)° generates a cone that is tangential to the detector (ro), thus an intensity is measured. **d-f.** The fiber orientation (α, β) = (70.5, 15)° generates a scattering cone that is not directed toward the detector. **g.** Simulated scattering amplitude profiles for different initial fiber orientations changing the simulated radii of the fibers, from left to right: 8, 100 and 150 nm. Note the different scales of the y-axis. **h**. Simulated scattering profiles for fiber bundles with varying fiber-orientation standard deviation parameters (σ) and the same initial main orientation. **i.** Probability density functions representing fiber orientation distributions for bundles centered at (α, β) = (90°, 0°), with different standard deviation values. Bounding box colors correspond to the dispersion parameters illustrated in panel (b). **j.** Schematic of the concept of fiber-orientation distribution.

**Extended Data Fig. 4.**
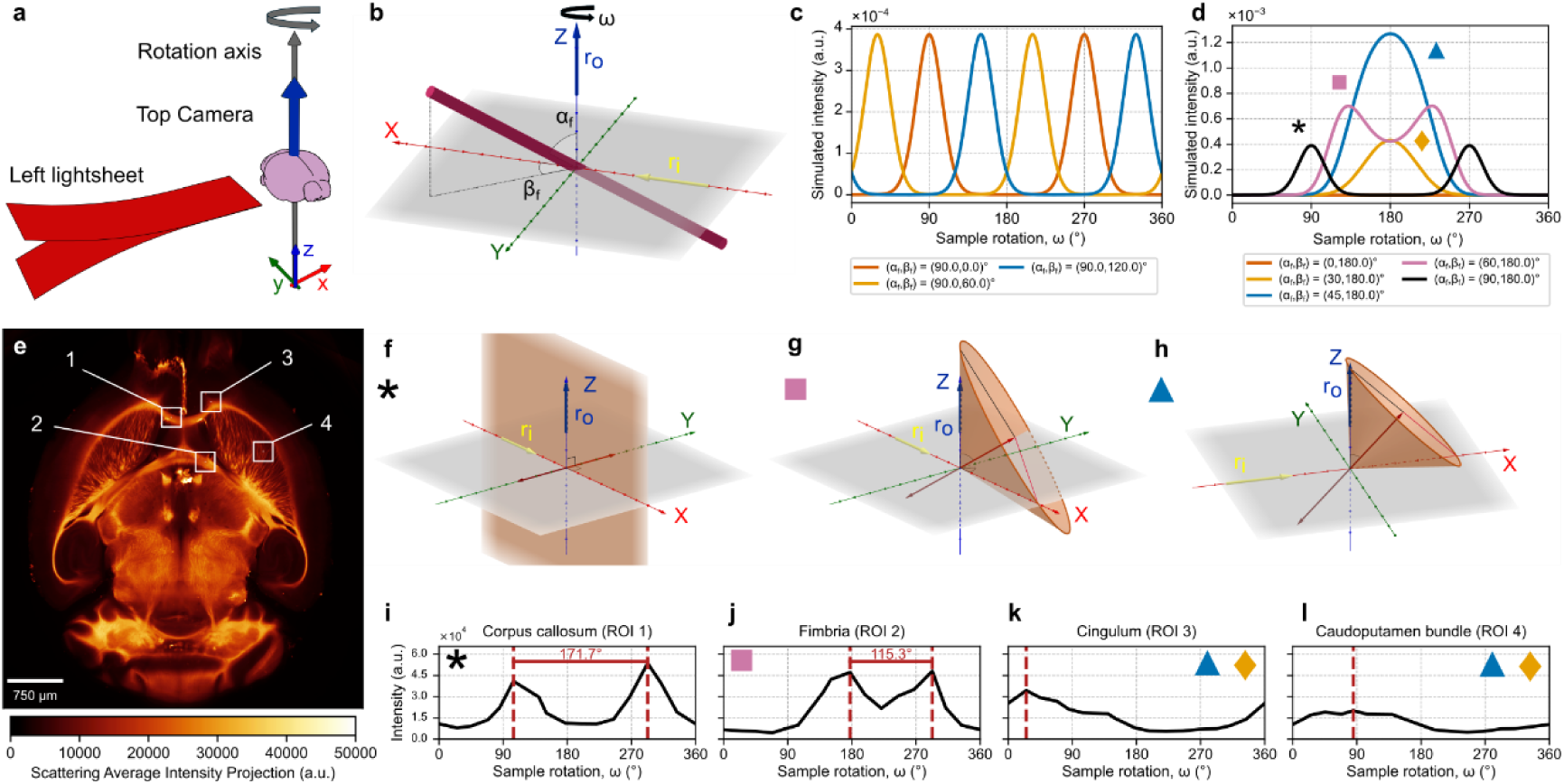
Simulated and experimental results through Z-axis rotation in an upright ultramicroscope system. **a.** Schematic of the ultramicroscope system. The detector is placed on the top of the sample, and the detector is aligned to the rotation axis. **b.** Parametrization of the 3D fiber orientation defined by tilt (αf) and azimuthal (βf) angles. ri and ro denote the light-sheet trajectory and detector position, respectively. **c.** Simulated scattering profiles for in-plane fibers (αf=90°) with varying initial azimuthal angle **d.** Simulated scattering profiles for fibers with the same initial azimuthal position (βf=180°) with varying tilt angle. The distance between the peaks depends on the tilt of the fibers, until they merge into a single peak for α = 45°. **e.** Average of the scattering signal across the 16 volumes and regions of interest. **f-h.** Schematic of the fiber position (red double arrow), and generated scattering cones (orange) in different positions where a peak intensity is measured. Note the different tilting of the fibers (**f**, α=90°, β=90°; **g**, α=60°, β≈54.74°; **h**, α=45°, β=0°). The top left icon links the schematics with the simulations (d). **i-l.** Mean scattering profiles in different regions of interest (6x6x1 voxel regions). Red dashed lines indicate the peak position and the distance between the peaks. Top icons link the scattering profile with a similar simulation.

**Extended Data Fig. 5.**
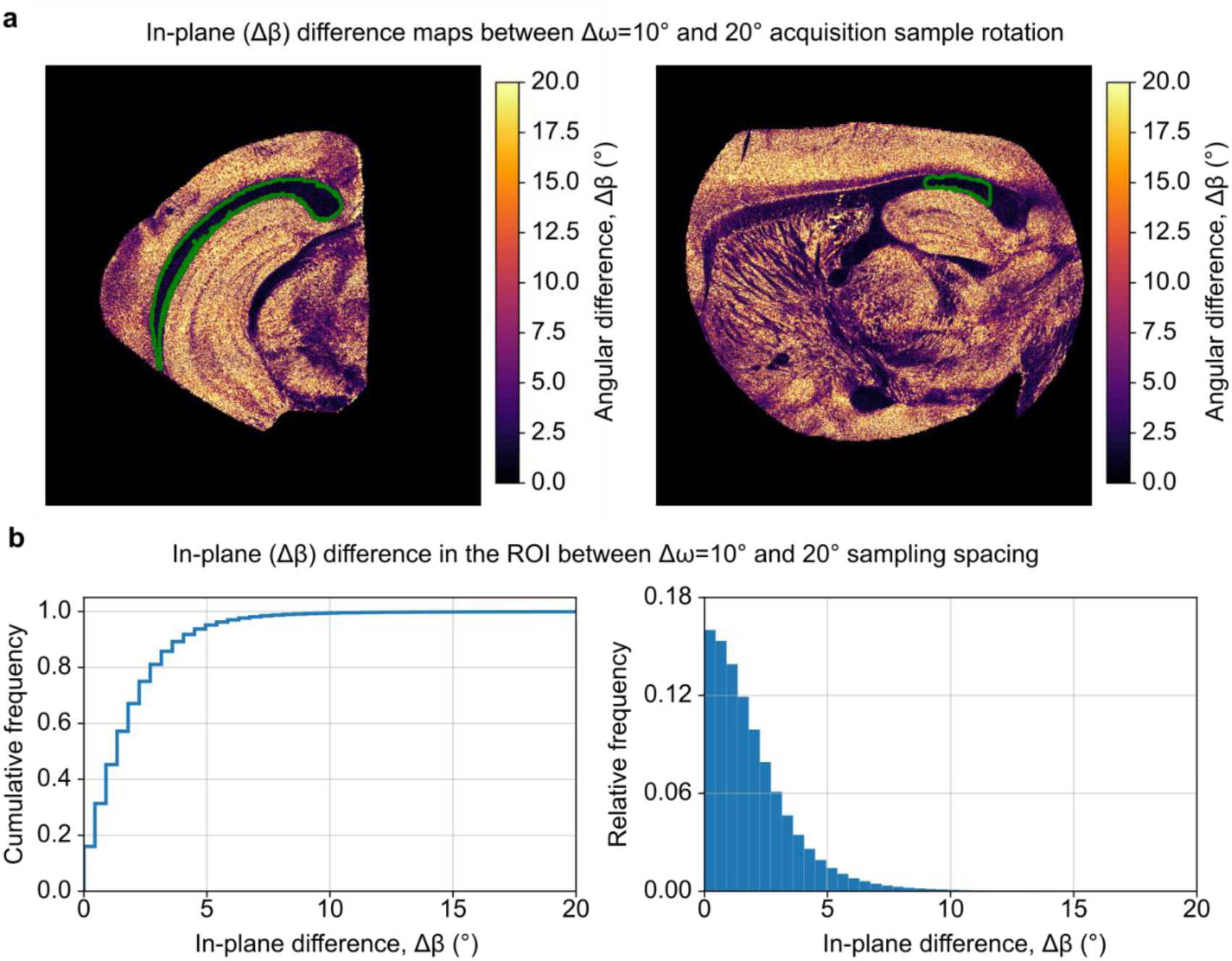
Experimental evaluation of the acquisition angular sampling step size on in-plane reconstruction. To evaluate the effect of varying angular sampling spacing in experimental data, we imaged a mouse brain hemisphere over a 360° range with 10° intervals (voxel size: 3.385×3.385×6.67 μm^3^). **a.** Angular difference maps the estimated in-plane orientation using all the mvLSSM acquired data (10° nominal angular sampling) and only every second volume (20° nominal angular sampling). In green, the region of interest for quantification. **b.** Cumulative (left) and relative (right) histograms of the in-plane angular difference in a mouse corpus callosum region segmented over 300 coronal slices. We obtained a mean ± standard deviation angular difference of 2.00 ± 2.06°.

**Extended Data Fig. 6.**
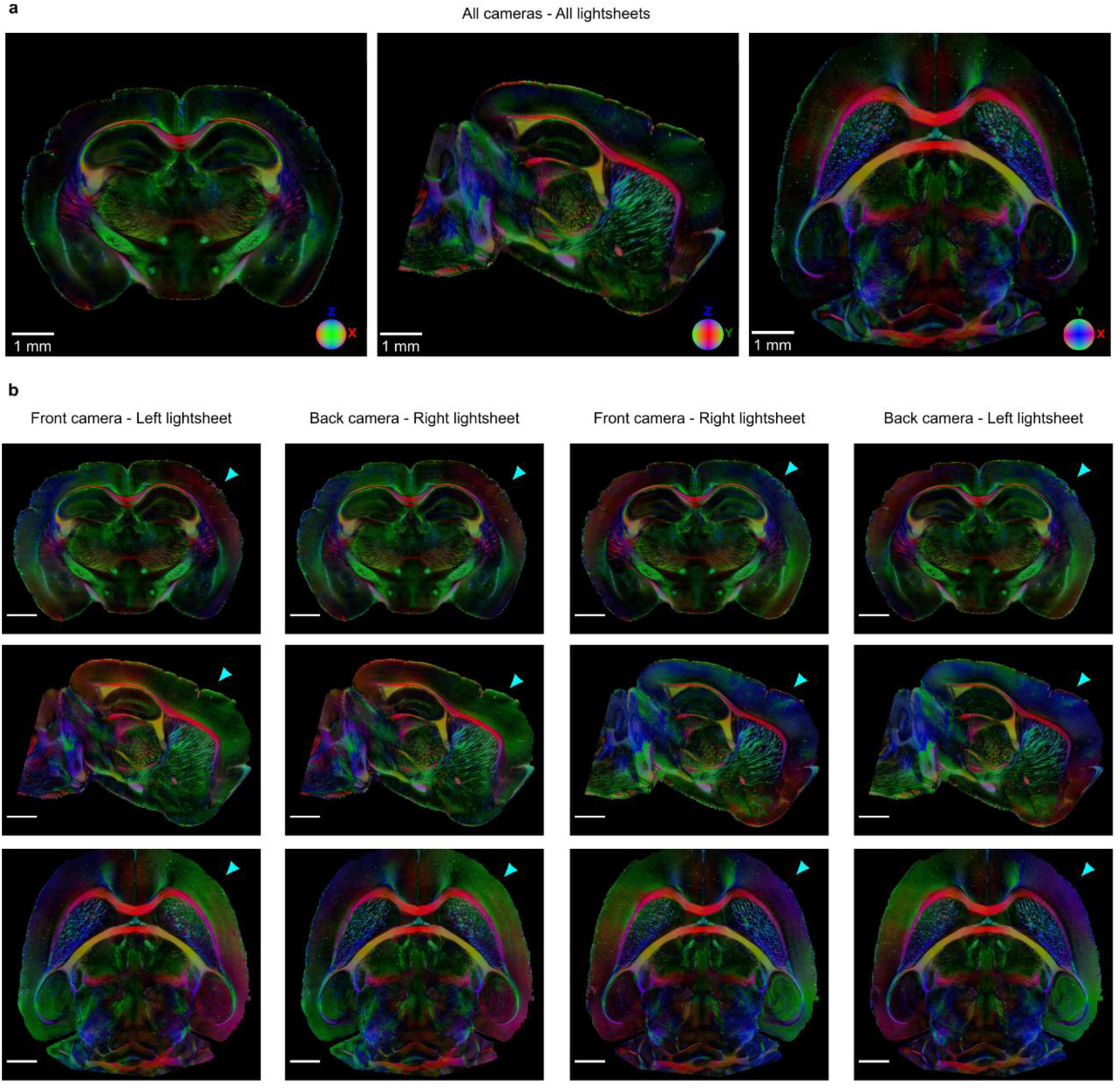
Comparison of tLSSM fitting using different camera/light-sheet pairs. **a**. Coronal, sagittal and axial views when using the four camera/light-sheet combinations to estimate the directions **b**. Coronal, sagittal and axial (row-wise) fitting for every camera/light-sheet pair (column-wise). Cyan arrowheads point to cortex areas where the estimated direction is different between fittings of opposing cameras or light-sheets, probably due to attenuation effects of light-sheet illumination across the sample, hence to the different optical paths for the peak position between views. **Scale:** 1 mm.

**Extended Data Fig. 7.**
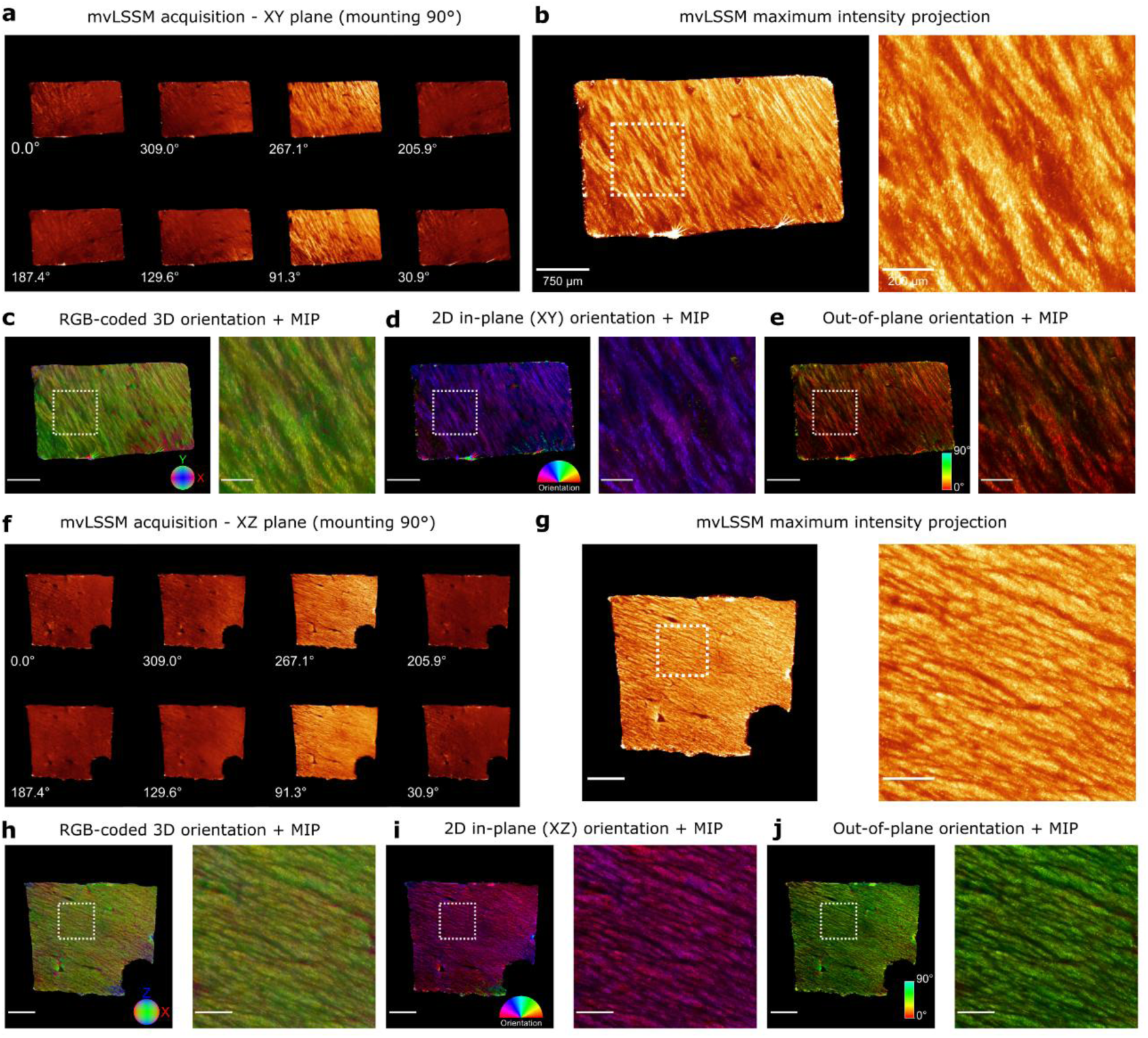
mvLSSM imaging and tLSSM of a chunk of the human corpus callosum. **a,f.** mvLSSM axial (**a**) and coronal (**f**) representative slices from selected views. **b, g.** Axial (**b**) and coronal (**g**) maximum intensity projection (MIP) of the mvLSSM acquisition. **c-e.** Representation of the estimated orientations in the axial slice overlaid onto the MIP as (**c**) RGB coded directions and decomposed into (**d**) in-plane XY orientation and (**e**) out-of-plane orientation, as the angle from the XY-plane towards the Z axis. **h-j.** Representation of the estimated orientations in the coronal slice overlaid onto the MIP as (**h**) RGB coded directions and decomposed into (**i**) in-plane XZ orientation and (**e**) out-of-plane orientation, as the angle from the XZ-plane towards the Y axis.

**Extended Data Fig. 8.**
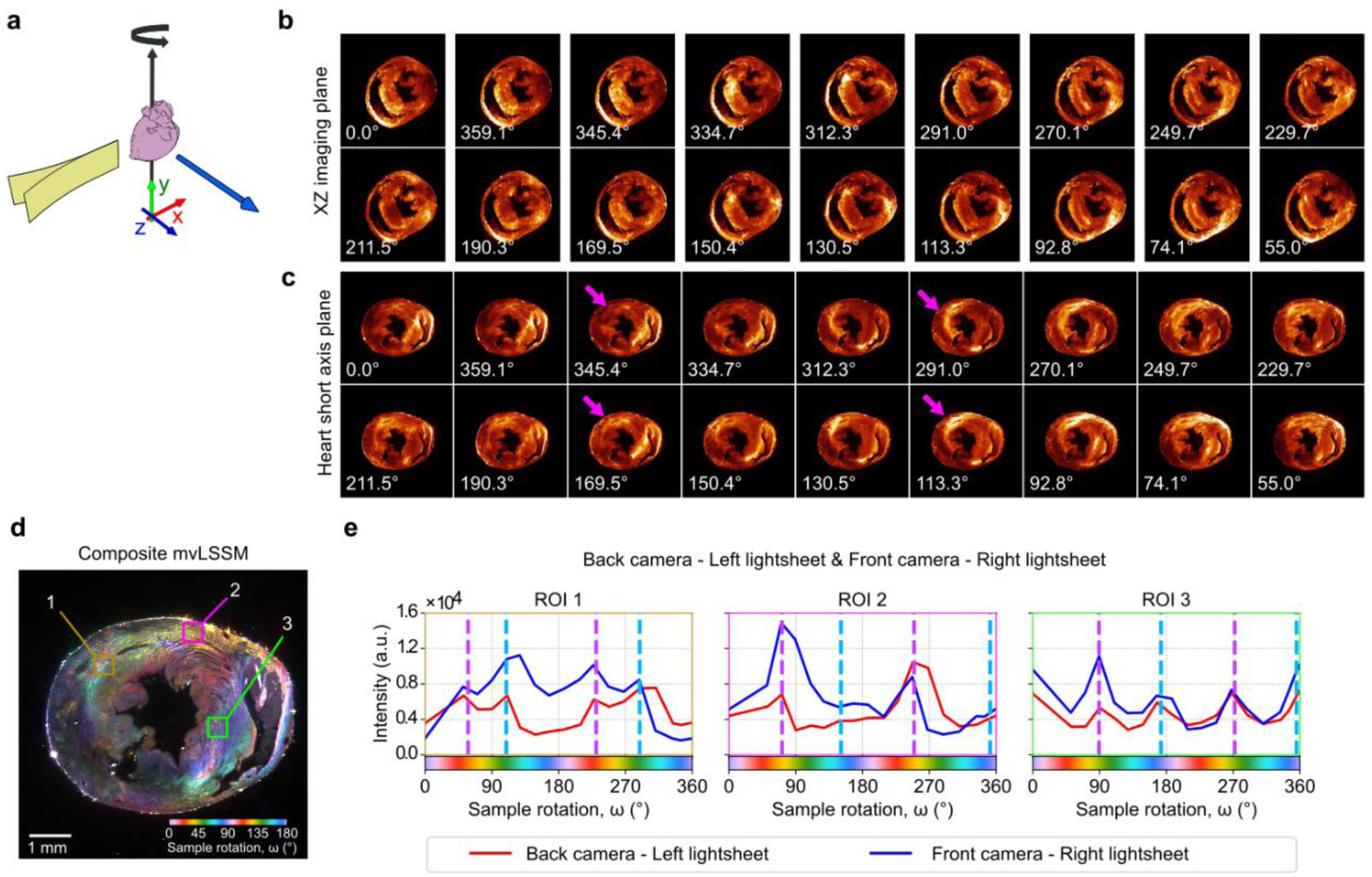
mvLSSM imaging of the mouse heart. **a-c.** mvLSSM acquisition of the mouse heart. To compensate for light-sheet degradation, LB and RF light-sheet/camera perspectives were merged through a maximum intensity projection. **a.** Schematics of the heart position with respect to the microscope. **b.** mvLSSM acquisition of an XZ slice in the imaging space (rotation axis perpendicular to the image). Magenta arrows point at an area with changing scattering intensity. **c.** mvLSSM of the same slice aligned to the heart short axis. **d.** Composite image showing the mvLSSM acquisition and 27×27×4.5 μm^3^ regions of interest. **e.** Averaged scattering profiles suggesting crossing fibers and estimated peak positions in the regions of interest.

