## Supplementary Information for "Mapping 3D tissue orientation with tensor-augmented light-sheet microscopy"

#### Supplementary Methods 1. Instrument characterization

##### Instrument for dual view light-sheet imaging of cleared samples and mvLSSM

The primary light-sheet instrument used and adapted in this study is a custom device based on macroscope detection, i.e. macroSPIM, with double sided detection (2× AZ100M macroscopes, Nikon, Japan) and double-sided illumination. Supplementary Figure 1 describes the optical layout with all components. Illumination arms: a laser combiner (LightHub, Omicron, Germany) feeds three solid-state lasers (488, 561, 638 nm) into an optical fiber, and the beams are first collimated with a parabolic fiber collimator (RC12APC01, Thorlabs Gmbh, Germany), then line- and power-selected with an acousto-optical tunable filter (AOTF 8 channels, AA-Optoelectronics, France). A half-wave plate (AHWP05M-600, Thorlabs) mounted on a motorized rotation stage (DDR25, Thorlabs) adjusts the linear polarization orientation before a 10x beam expander (Sill Optics, Germany) collimates the beams into a >20 mm diameter. A manual slit enables to control the aperture of the beam to tune the light-sheet profile (i.e. thickness) and a large galvo mirror (GVS211 “Y-axis”, Thorlabs) enables one to distribute the beams onto the desired illumination arm. Light-sheet shaping entails on each side: an electrotunable lens (ETL16, Optotune) to control lateral light-sheet waist position, a one-axis galvo (GVS211, Thorlabs) for pivot scanning in Y to eliminate striping[1] and reduce speckled noise in scattered images (as shown previously by Di Battista et al.[2]), a relay 4f telescope (f1 = f2 = 160 mm) and a 50 mm achromatic doublet cylindrical lens (ACY254-050-A, Thorlabs). In detection, each side entails: a macroscope mounted on a linear motor (L511, Physik Instrumente, Germany) for focus control that also holds a filterwheel (Lambda 10-B, Sutter Instrument, CA, USA) and a Flash4.0v2 sCMOS camera (Hamamatsu, Japan). The sample is mounted vertically in a quartz cuvette (20×20×35 mm, 2 mm quartz wall) and a XYZ+rotation bracket (motors L505, rotation motor M116DGH, Physik Instrumente, Germany). The sample rotation is contact-free and made with a custom magnetic rotation mechanism, where rotating a quadripolar “master” magnet stack under the cuvette acts on a “slave” magnet inside the cuvette (custom magnets from Alga Magneti SRL, Italy). To limit frictions, a radial bearing is placed under the slave magnet, and a silicon pad is shaped to fit onto it to enable inserting pins (100microns Austerlitz insect pins, Entomoravia, Czech Republic) that will hold the agarose block in place. Rotation accuracy of the assembly was estimated to be below +/- 3° around the target position, explaining why rotation steps have intrinsic variability, due to non-contact magnetic movements, that are considered in all aspects of the multiview data processing.

#### Static vs. ASLM light-sheet formation

As depicted in Supplementary Fig. 1-e, light-sheet imaging can be performed with static illumination, shaped to cover the entire field of view (FOV) with minimal thickening of the illumination sheet on its edges. For a magnification of 1.44× and a FOV of 9.24×9.24 mm, used for full brain imaging, LS waist is typically below 20um in the center and below 25 µm on the edges. Static LS was used to generate images shown in Fig. 1 and Extended Data Fig. 1 of this paper. Alternatively, axially swept light-sheet microscopy (ASLM)[3] can be enabled by opening the input beam to maximum aperture and rastering the thinnest LS waist to reconstruct an image made of, typically 13, bands of highest axial resolution (i.e. thinnest light-sheet thickness). Our system cannot make use of rolling shutter, hence the reconstruction of ASLM images requires the acquisition of individual images, reconstructed at the acquisition with the custom control software (programmed in Labview 2016, National Instruments, Texas, USA: code available upon request). With a >10 mm diameter beam and a 50 mm cylindrical lens focusing through about 1 cm of imaging medium (BABB or DBE, refractive index around 1.56), the resulting LS waist is below 6 µm. This includes the pivot movement which is estimated to thicken the LS by about 20% compared to non-pivoted beam due to off-axis alignment effects. For most datasets presented here, volume imaging was performed with z µm steps below the half width of LSw, hence ensuring proper and continuous axial sampling (see Supplementary Table 1).

**Resolution and acquisition strategy across the datasets**

Several acquisition strategies were employed depending on the purpose: static light-sheet (LS) provides milder optical sectioning for axially subsampled imaging aimed to visualize with coarse resolution, while ASLM is required to ensure the best axial resolution for mvLSSM. Supplementary Table 1 reports the main settings used for each sample and the figures where they are displayed.

**Supplementary Table 1. Acquisition parameters for each dataset.**

| Sample, fixation, clearing | Shown in Figures | Magnification / estimated Numerical Aperture (NA) | XY pixel size / Z step (µm) | Estimated LS waist in the center of the field-of-view (µm) | Single view or mvLSSM (Multiview steps in °), Use of ASLM (number of bands) or static LS |
|---|---|---|---|---|---|
| Mouse brain #1 unlabelled, PFA, iDISCO+ | Fig. 1. Extended data Fig. 1 | 1.44× NA ≈ 0.07 | 4.514/ 4.514 | <20 | Single view, Static LS |
| Mouse brain #2, MBP647, PFA, Labelled, iDISCO+ | Fig. 1, 5a-c Extended data Fig. 1 | 1.44× NA ≈ 0.07 | 4.514/ 4.514 | <20 | Single view and mvLSSM (20° steps), Static LS |
| Mouse brain #3, unlabelled, fixed in Glioxal, iDISCO+ | Extended data Fig. 1aa, ab, af, ag | 4.8× NA ≈ 0.153 | 1.35/50 | <10 | Single view, Static LS |
| Mouse brain #4, unlabelled, PFA, iDISCO+ | Extended data Fig.1y, z, ad, ae | 4.8× NA ≈ 0.153 | 1.35/50 | <10 | Single view, Static LS |
| Mouse brain #1, unlabelled, PFA iDISCO+ | Fig. 2 (all), 3, 4 (all), Extended Data Figs. 2, 7 | 1.44× NA ≈ 0.07 | 4.514/ 4.514 | <10 | mvLSSM (20° steps) ASLM (11) |
| Mouse brain hemisphere #5, unlabelled, PFA iDISCO+ | Extended Data Fig. 6 | 1.44× NA ≈ 0.07 | 3.385/ 6.670 | <20 | mvLSSM (10° steps), Static LS |
| Mouse brain #6, unlabelled, PFA iDISCO+ | Extended Data Fig. 5 | 1.26× NA ≈ 0.5 | 5.160/ 10 | <20 | mvLSSM (22.5° steps around Z), Static LS |
| Heart (mouse), PFA, iDISCO+ | Extended Data Fig. 9 | 1.44× NA ≈ 0.07 | 4.514/ 4.514 | <6 | mvLSSM (20° steps), ASLM (11) |
| Human brain, (chunk), formalin, iDISCO + | Fig. 5, Extended Data Fig. 8 | 1.44× NA ≈ 0.07 | 2.462/ 2.462 | <8 | mvLSSM (20° steps), ASLM (9) |
| Horse sciatic nerve #1, PFA, Me:OH+ BABB | Suppl. Fig. 2 | 1.92× NA ≈ 0.095 | 3.385/ 3.385 | <20 | mvLSSM (20° steps), Static LS |
| Horse sciatic nerve #2, PFA, CUBIC | Suppl. Fig. 3, 4 | 1.92× NA ≈ 0.095 | 3.385/ 3.385 | <20 | mvLSSM (20° steps), Static LS |

#### Supplementary Note 1. Polarization dependence of LSSM signal

We characterized the scattering signal dependence on light polarization in mouse brains, as shown in Supplementary Fig. 1f–p. A motorized rotation mount controlling a half-wave plate was used to adjust the linear polarization of the light-sheet, ensuring the same polarization orientation for both left and right light-sheets. Supplementary Fig. 1f-k shows that switching to p-polarization (p-pol: perpendicular to the light-sheet plane, see schematic in Supplementary Fig. 1n) from s-polarization (s-pol: parallel to the light-sheet) results in switching from a maximized scattered signal to a minimum signal (imaging at 1.44× magnification). Interestingly, the same effect is observed for both light-sheets at the same time (compare Supplementary Fig. 1g vs. 1j for the left light-sheet and 1i to 1k for the right).

To further investigate this polarization-dependent effect, particularly the sensitivity of fiber-associated signals relative to background or non-fiber-like scatterers, we imaged a hemibrain at 4.8× magnification. LSSM volumes were acquired at 20° polarization rotation intervals. Supplementary Fig. 1l-m displays merged views (front and back detection) centered on the caudoputamen and corpus callosum, showing strong LSSM fibers signal (regions F1-F4). Supplementary Fig. 1m also shows an area close to the hemibrain dissection plane (antero-posterior cut), where the cut-exposed surface and a large vessel are visible and covered with apparent point-like scatterers (e.g. point P3). By comparing the specific LSSM signal variations upon polarization rotation in fiber-like regions (F1- F4), cortex (C1) and non-fiber-like regions (P1, P2, P3), we could observe and measure that: (i) all regions show some intensity modulation concomitant with light polarization rotation, but in different amount/amplitude, and show a redundancy or periodicity of the signal (i.e. minimum and maximum peaks returning) upon rotation with 180° period; (ii) fiber-specific regions (F1-F4) and cortex (C1) modulate with much greater and in-phase amplitude, reaching almost complete extinction (below 4-5% of maximum intensity at s-pol) at p-pol. In contrast, background regions (outside the sample, 30% loss at p-pol) and non-fibers scatterers (P1-P3 down to 40-50% loss at p-pol) show only partial, and out-of-phase modulation.

From these observations, we conclude that the scattered signal originating from axonal bundles is completely polarization-dependent and can be sent out of the detection axis (cameras) by rotating polarization. This phenomenon is reminiscent of Rayleigh scattering detection that, as shown previously[2,4], is present in biologically cleared samples. Rayleigh scattering is assumed to stem from subwavelength- or submicron-sized tissue components and can be “extinguished” (for the observer) in the same polarization-dependent manner, i.e. by rotating polarization it can be sent out of the detection point. In Di Battista et al. (2019)[2], this was shown in agarose (i.e. small particles/mesh smaller than wavelength) and Simó et al. (2024)[4] this was used to eliminate Rayleigh scattering from agarose and tumor tissues (i.e. very small subwavelength debris). The observation of milder LSSM modulation on point-like particles at the brain surface has two possible explanations: (i) the polarization rotation affects the light-sheet focusing accuracy, hence both the intensity and focus position may vary locally and induce variations; (ii) the scatterers are larger and can respond to a different, possibly Mie-scattering like, regime that is less polarization dependent and scatters light more isotropically. This interpretation is directly supported by Supplementary Fig. 1l-m where many point-like scatterers contribute equally to both cameras’ images (see white spots in Supplementary Fig. 1m, merges of green and magenta).

In conclusion, we have related fiber scattering detection to the Rayleigh scattering regime based on polarization behavior, which also suggests that fiber-dependent scattering arises from a sub-wavelength light-to-tissue mechanism that may be induced by subwavelength fibers organization or structure, although this study does not demonstrate it directly.

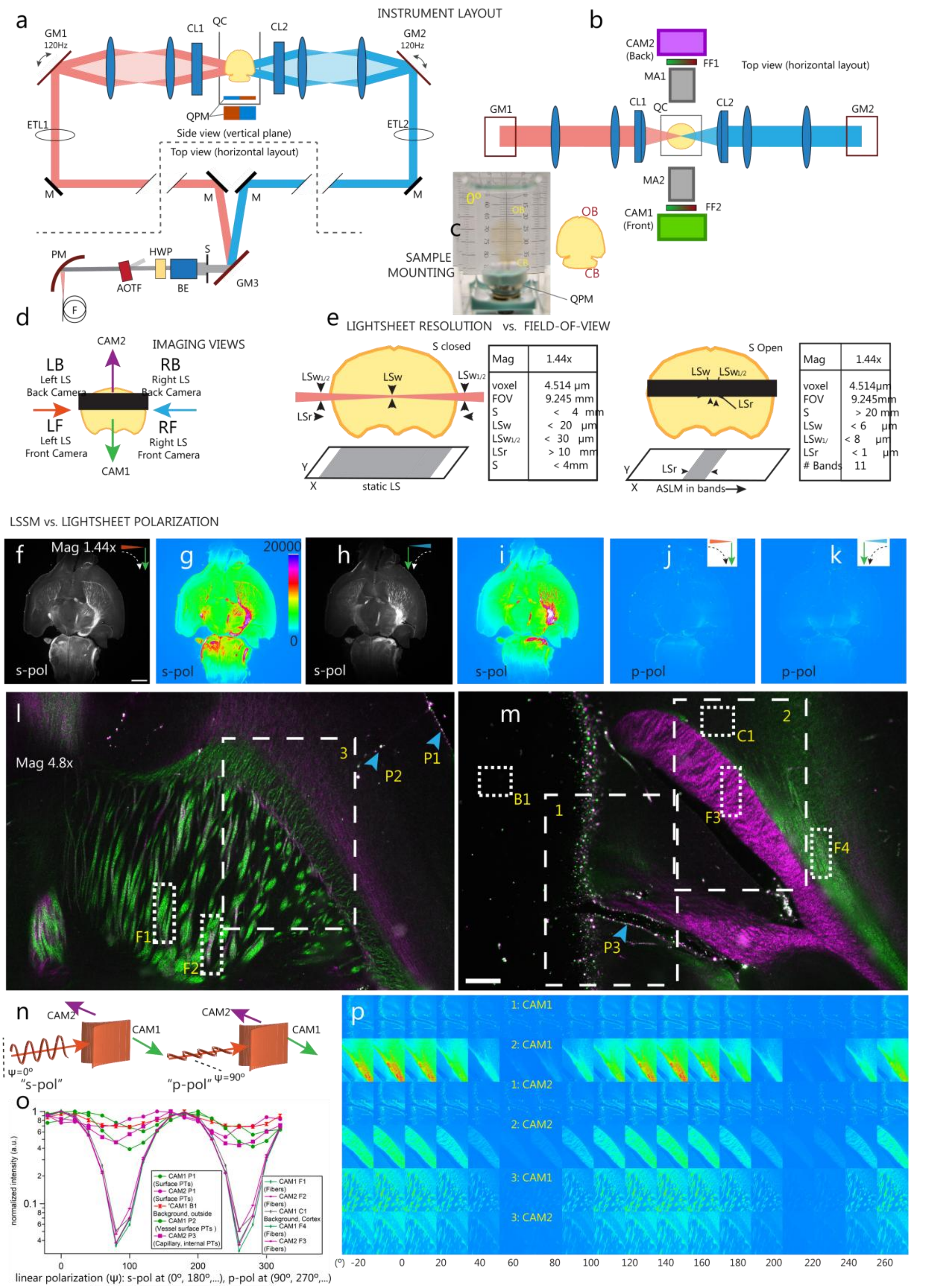


**Supplementary Figure 1: LSSM instrument and polarization dependence of LSSM signal. a.** side view (top) and top view (bottom) of illumination arms: laser beams are collimated (PM), power-selected (AOTF), linear polarization is turned (HWP) and further expanded (BE). Light-sheet (LS) aperture is controlled (S) and illumination side is steered (GM3). Beam divergence is tuned

(ETL1,2). Pivot scanning at 120Hz (GM1,2) before relay telescopes and light-sheet formation (CL1,2). F: optical fiber, AOTF: acousto-optical tunable filter, HWP: Half wave plate, BE: 10× Beam expander, S: Manual slit aperture, M: Mirror, ETL: Electrotunable lens. GM: Galvo mirror. CL: Cylindrical lens 50 mm. Samples sit inside a quartz cuvette (QC), on a magnetic base (QPM: quadripolar magnet) and rotate through the action of a QPM magnet stack below QC. **b.** Top view of illumination and detection arms: motorized macroscope units (MA1,2) on each side with fluorescence filters (FF1,2) wheels, and sCMOS cameras (CAM1,2). **c.** Magnet based inside cuvette made of bearing (below magnet) and silicon pad (above magnet). Sample stands on insect pins. Shown sample: cleared mouse brain (CB: cerebellum, OB: Olfactory bulb). **d.** Four imaging views combining pairs of Illumination and detection sides: LF, LB, RF, RB (L: Left for CAM1, R: Right, Front: CAM1, Back: CAM2). **e.** Representation (not at scale with the sample drawing) of LS shaping relative to field of view (FOV). With static LS, a single image is captured with LSw and LSw1/2 thicknesses (w, for waist) at the center and edge of FOV respectively, formed with closed slit, for an effective light-sheet Rayleigh length Lr. In ASLM mode, the slit is open and LS focused on a thinner waist with shorted Lr. The ASLM image is formed by rastering LS along the x-axis and acquiring bands (typically 11 or 13 bands), recombined into a single image. **d-e** are top views.

**f-p.** linear polarization dependence. Scattered signal is maximized on both cameras/side with polarization perpendicular to light-sheet/image plane (p-pol: parallel to incidence plane), and minimum when parallel to light-sheet (s-pol: perpendicular to incidence plane). **f,g.** single plane with LF view and p-pol, grayscale and color-coded (intensity scale 0 to 20000), **h,i.** RF view, Greyscale and color. **j.** LF perspective with s-pol, **k.** s-pol with RF view. **l,m.** merged LF (green) and LB (magenta) in two different planes. F1,2,3,4: Fibers regions for intensities reported in o; P1,2,3: non-fiber scattering objects (e.g. dust) at the brain surface (P1), inside a capillary in the cortex (P2), and at vessel inner surface (P3). B1: background region outside the hemibrain. **n.** Schematic view of p-pol and s-pol for LF and LB. **o.** Scattered Intensity plot (log scale) upon polarization orientation in regions F1-4, P1-3 and B1. The graph shows that fiber-specific signals have a much higher polarization-dependence with deeper modulation than background or non-fiber scattering objects. **p**. Polarization is turned in steps of 20°, intensity of dashed regions 1,2,3 from **l-m.** for LF (CAM1) and LB (CAM2), is shown in color scale (intensity scale 0 to 20000 in **g**). Whole mouse brain imaged at 1.44× magnification in **f-k.** and a hemibrain from a different animal in 4.8x in **l, m, p**. Scale: in **f.** 1mm, **m.** 0.2mm.

### Supplementary Note 2. Sciatic nerve analysis

#### mvLSSM on sciatic nerve and analysis of fiber orientation to LSSM signal detection.

A horse sciatic nerve sample was cleared with two different clearing protocols (see Supplementary Methods 5) and imaged by mvLSSM to characterize the orientation dependence of the LSSM signal. The nerve fibers in this sample were considered macroscopically straight and aligned, hence of known or easy to retrieve orientation at the macroscopic scale (i.e. over the length of the fascicles, a few millimeters). For both samples, mvLSSM was achieved by rotating the sample in steps of about 20° across the full 360° range (18 views), and volume images were acquired at isotropic lateral-to-axial sampling, at a magnification of 1.92x (voxel size $3.385^3$ $\mu m^3$), ensuring the entire sample remained within the field of view throughout rotation. In all Supplementary Fig. 2-4, the rotation axis for mvLSSM is denoted along the Y-axis, perpendicular to the light-sheet-to-camera plane.

#### BABB-based clearing

With solvent based clearing (BABB; Supplementary Fig. 2) we imaged a first chunk of nerve in two different orientations: with fiber fascicles oriented in the plane light-sheet-to-camera and rotating (Supplementary Fig. 2a-c); and with fibers aligned along the rotation axis Y (Supplementary Fig. 2w). We first observed pronounced differences between autofluorescence and scattering. Across all views, fascicles exhibited modulated LSSM signals, whereas autofluorescence revealed stable intensity in connective tissues that were not visible in scattering (Supplementary Fig. 2e–f, i–j; multiview autofluorescence images not shown). These findings suggest that nerve fibers are the main source of scattered signal while cleared connective tissues do not contribute to it.

Notably, the scattered signal from individual fascicles displayed periodic undulating patterns along the fiber main axis direction, with intensity peaks occurring in two distinct sets of views approximately 180° apart (Supplementary Fig. 2d; compare views a6–a9 with a15–a18). This 180° redundancy is consistent with the geometric symmetry of the fibers with respect to the microscope elements, which return to be aligned along the same direction for this specific sample positioning and rotation. By combining views into rotation-encoded color projections (Supplementary Fig. 2m-v), we show that the patterns intercalate in successive views, i.e. the periodic patterns shift or “slide” along the fiber axis from one view to the next, or in other words patterns are complementary between views. We also observed (see grey scale images) that internal fibers in the fascicles are distinguishable, and curl or undulate in central regions. Looking at cross sections (Supplementary Fig. 2r-v), we could resolve the color distribution and observe that patterns are preferentially not centric/radial.

In the second configuration, where fibers were aligned with the rotation axis, the mvLSSM scattered signal remained stable across all views, with some local patterning yet of much less obvious periodicity. Rotation color-encoded image (Supplementary Fig. 2z, ac) showed scrambled colors from the full color scale and no specific dominance from a specific view.

From these observations, we conclude that (i) straight nerve fibers scatter light orientationally, given they modulate the collected scattered light and are the only moving parts in these experiments (camera and light-sheet are static); (ii) there is a 180° symmetry in the maximum scattering by fibers consistent with the geometric symmetry of the fibers that return to the same orientation upon 180° rotation; (iii) there is no LSSM modulation when fibers and rotation axis are colinear, reinforcing the idea that LSSM is steered by fiber orientation which does not change in this specific configuration across the views; and (iv) although macroscopically the fibers provide a global response (i.e. the whole fiber lights up) to rotation, at the microscopic level a periodic pattern exists that suggests there is a possible relation between the local internal fibers orientation, inside the fascicle where curling is visible, and the maximum scattered signal detected. If so, this would suggest that nerve fascicles scatter light with fair angular sharpness and that mvLSSM is accurate and/or sensitive enough to capture local angular variations of the fibers, hence establishing a direct and quantitative relation between fiber orientation and multiview LSSM signal collection. In the next figures (Supplementary Fig. 3 and 4), we unravel a quantitative analysis to address this hypothesis with a second clearing protocol on a new chunk of the same nerve sample.

#### CUBIC-based clearing

Supplementary Fig. 3g shows a mvLSSM acquisition scheme on a CUBIC-cleared chunk, rotated with nerve fibers in the light-sheet-to-detection plane (like in Supplementary Fig. 2c). Consistent with prior observations in the BABB-cleared sample, we observed strong differences between an expected rotation-invariant autofluorescence distribution (autofluorescence rotated images not shown) and a two-peaked LSSM signal modulation (Supplementary Fig. 3a-f, all views in Fig. 3h). Additionally, we observed differential LSSM responses between the two detection cameras (Supplementary Fig. 3d vs 3e), similar to effects previously noted in brain tissue (Fig. 1e,g).

Interestingly, rotation color-encoding of a registered fascicle showed, again, a periodic pattern of LSSM signal maxima along the axis of the fibers (Supplementary Fig. 3l, 3n), consistent with the BABB-cleared sample. Curling of individual fibers was also evident (Supplementary Fig. 3k, 3m, 3o). Altogether, these observations show that nerve fibers' light scattering in sciatic fascicles is conserved in two very different clearing protocols chemically (solvent-based vs. water based), suggesting that nerve fibers interact with light in a robust and reproducible manner.

Noteworthy, two differences were observed this time with CUBIC clearing. Firstly, connective tissues exhibited a strong LSSM response, in particular in the perineurum (see arcs of high intensity around the fascicle section in Supplementary Fig. 3i,j). Though we ignore this source of scattering in the present analysis, we can hypothesize that perineurum fibers, known to organize circularly around fascicles[5], could be responsible for this signal. The absence of this signal in the solvent-cleared sample suggests that water-based clearing may better preserve this tissue component. Secondly, we observed that internal fibers were more distinguishable than with BABB clearing (compare, at the same magnification, Supplementary Fig. 2m-p with Supplementary Fig. 3k,m,o, and cross sections Supplementary Fig. 2r with Supplementary Fig. 3i,j,q,t). Indeed, cross sections of the fascicle (Supplementary Fig. 3t,q) show ring-like structures (see pink/red arrowheads), reminiscent of myelinated rings which are prominent in horse sciatic fascicles[5]. Noteworthy, such rings are not visible on all views, as shown by comparing two consecutive angle views (rings in 3q are not visible in 3p, rings in 3t not visible in 3u).

Our interpretation for this observation is two-fold. (i) The visibility of ring-like structures in the CUBIC-cleared sample, but not in the BABB-cleared sample, may result from differential clearing-induced side effects. CUBIC is known to cause mild swelling, whereas MeOH:BABB induces mild shrinkage. At the moderate imaging magnification used (1.92×), myelinated rings may appear larger in CUBIC samples, hence better optically resolved. Furthermore, (ii) optical resolution is not isotropic in such a dataset (i.e. xy resolution approximately 3-fold better than axial). Therefore, it is expected that ring-like structures, at the limit of resolution power at 1.92× magnification, appear only in those rotated views where fibers are aligned with the optical axis of the microscope, whereas other views would feature axial resolution at a large angle with fibers, hence introducing blurring. This is the case with the selected view in Supplementary Fig. 3i-w: views a9 and a10 correspond to angles around 180° between light-sheet and fiber axis, hence the better lateral resolution is perpendicular to fibers and they get better resolved.

Altogether, this dataset (Supplementary Fig. 3) shows that in a selected angular view, imaging resolution is sufficient to visualize fascicles' fibers. Qualitative observation points suggest a correlation between fiber orientation and the angular view associated with maximum LSSM signal intensity. This is illustrated by the color (i.e. angular view) dominance in Supplementary Fig. 3s (red color) with fibers pointing "up-left", and Supplementary Fig. 3v (cyan-green color) pointing "up-right".

#### Quantitative mapping of fiber orientation and scattering peak displacement

To establish a quantitative relationship between local fiber orientation and LSSM scattering peak positions, we developed an analysis pipeline that operates at the voxel level within a 3D section of the fascicle shown in Supplementary Fig. 3. This pipeline quantifies, for each voxel, both the angular position of scattering peaks and the local orientation of fascicle fibers.

Using the registered angular views, we selected a fascicle fragment (an XZ plane is shown in Supplementary Fig. 4i–m) and manually segmented its boundaries. then subdivided it into a 2D grid of 10×10 pixel boxes (i.e. 34×34 µm), for each plane across the Y-axis. We also measured the fascicle center (FC on Supplementary Fig. 4m), by fitting an ellipse (not shown) to the fascicle cross-section, allowing the calculation of the radial position (R) of each grid box relative to the center. In each box, we measured the mvLSSM intensities and fitted the profile into a double gaussian plot (Supplementary Fig. 4b) to retrieve the angular position of the two peaks referred to as $\tau_A$ and $\tau_B$. Then, local fiber orientation was then estimated using the Fiji plugin "OrientationJ"[6], which computes the gradient structure tensor within a defined neighborhood. Measurements were performed in 40×20 pixel rectangles centered on each grid box (40 pixels perpendicular and 20 pixels along to the fiber axis). The angle between the estimated 2D direction and the fiber axis (vertical axis on the images Supp. Fig 4d-p), yielded an angular distribution (or histogram, centered on 0° for the fiber axis in Supplementary Fig. 4c). The distribution of angles features a peak, whose center is captured by fitting a Gaussian function, denoting the peak angle as **Beta**. Supplementary Fig. 4n,o,p shows example distributions of $\tau_A$, $\tau_B$-180°, and Beta in a selected XZ plane, where the pattern distribution shows strong similarities between $\tau_A$ and $\tau_B$-180°, and milder graphical similarity between Beta and the two formers.

To further evaluate these similarities, we plot heatmaps (Supplementary Fig. 4q-v) for $\tau_A$, $\tau_B$ -180°, Beta, the summed full-width at half-maximum (FWHM) of the two peaks ([c+f], see equation in Supplementary Fig. 4b), the ratio $\tau_A/( \tau_B -180°)$, and the difference $\tau_B - \tau_A$. These maps are plotted along the fiber axis (Y, bottom axes) and radial position (R, left axes), representing a 3D dataset projected onto a 2D image. The heatmaps reveal the following: Firstly, (i) the correlation between $\tau_A$ and $\tau_B$ is strongly confirmed visually, indicating that the angular displacement of the LSSM scattering peaks occurs in a coordinated manner across the fascicle (Supplementary Fig. 4r,s). This observation is further supported by the homogeneity of the ratio and difference maps, with the latter consistently centered around 180°, as anticipated (Supplementary Fig. 4u,v). Supplementary Fig. 4t provides insight into the variability of the measured data by displaying the sum of the two peaks' FWHM values ([c+f]), which highlights regions with sharper scattering profiles (i.e. sharper scattering profiles, in green). These regions correspond to areas in Supplementary Fig. 4u–v where the $\tau_A/(\tau_B\text{-}180°)$ ratio approaches 1, and the difference ($\tau_B - \tau_A$) is closer to 180°, reinforcing the interpretation that fiber scattering is symmetric and exhibits angular redundancy. We also plot all data points of $\tau_A$ vs. [$\tau_B$-180°] in Supplementary Fig. 4w and show that filtering the data to plot only those points standing within the FWHM (i.e. removing outliers, Supplementary Fig. 4x) increases their visual linear relation (see linear fit slope increase from 4w to 4x).

Secondly, (ii) a milder similarity between $\zeta$ and both $\tau_A$ and $\tau_B$ can also be graphically appreciated (Supplementary Fig. 4q to compare with -r and -s), suggesting a relation between fiber orientation and scattering peaks displacements throughout the body of the fascicle. Similar to Supplementary Fig. 4x, we plot $\zeta$ against $\tau_A$ for the most significant LSSM peaks (i.e. removing large peaks outliers of [c] and [f]). The resulting scatter plot reveals a distinguishable linear trend, supporting the hypothesis of a link between local fiber orientation and scattering behavior (Supplementary Fig. 4y).

Altogether, despite the limited resolution (i.e. imaging at 1.92× magnification), limited clearing efficiency and limited accuracy of volume registration (due to the latter two points), we believe we demonstrated that a clear and robust relationship between fiber orientation and LSSM scattering in the specific configuration where nerve fibers are oriented perpendicular to the rotation axis. This brings evidence to support the interpretation of scattering peaks redundancy, shown in the main text of this manuscript in a more complex sample such as brain neural pathways, and to support the choice of the geometrical cylinder scattering model to interpret orientation retrieval in tLSSM.

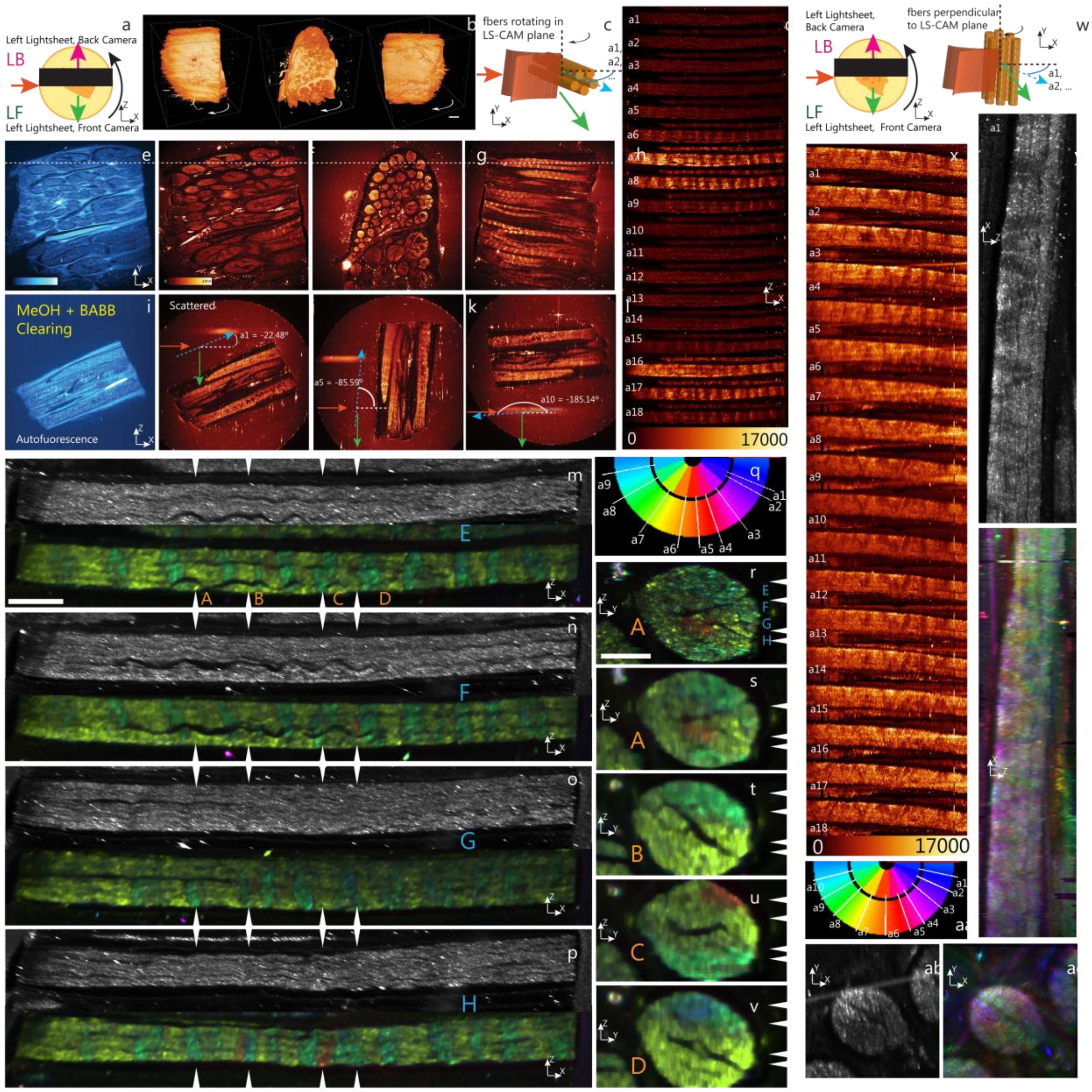


**Supplementary Figure 2**: **Orientation-dependent scattering in BABB-cleared sciatic nerve fibers.** **a-v**: The sciatic nerve fascicles, cleared in BABB, rotate along the light-sheet-to-camera plane. **a**. Top schematic view of sample and rotation orientation. **b**. 3D rendering of the sample at three selected rotations/views. **c**. 3D schematic relation of light-sheet, fiber fascicles and rotation axis. **d.** single XZ plane of aligned/registered views of one selected fiber bundle, for all angles a1, ..., a18 (20° average step between views). Several views (i.e. a6 to a9 and a15 to a18) clearly show a LSSM signal increase, an internal discontinuous patchy pattern along the fascicles and a half-turn redundancy (e.g. approximately 180° between a9 and a16). Bottom: color scale, intensity levels. **e-h.** XY views, color scale in e and f. **i-l.** XZ views from dashed line in **e-h**, color scale in F. **e,i.** Autofluorescence; **f-h,j-l.** LSSM signal showing parallel fascicles in the nerve chunk. Background signal around the sample stems from scattering by the cylindrical agarose block, visible in J-L. **m-p.** four selected XZ planes (at different Y heights) across the fascicle show (top) greyscale intensity of view a13 (of minimal signal from D) and (bottom) the same image color-coded in rotation with signals from views a1 to a18 (see Supplementary Note 2 for description of color-coding of the panels). **q.** Relation between color code ("BIOP12colors.lut" in Fiji) and angles a1 to a9, showing the origin of red-green-cyan dominance from views a6 to a9. **r-v:** same approach, but for selected YZ cross sections at different X positions along the fiber. White arrowheads show corresponding plane positions from XZ to YZ images, labelled (A) to (D) and (E) to (H). Image in R shows raw data (colored) while images in **m-p** and **s-v** use a filtered image (removing outliers) to ease the distinction of color patterns. Note that images M to P show local buckling of the fibers inside the fascicle and possibly a wider angular distribution of the signal with cyan and red colors (as opposed to green) in the fiber center (as opposed to the fiber tips). While in XZ images, color patterns seem periodic along the fiber axis, YZ images show that the angle-dependent distribution of scattering is more complex. **w-ac.** The sciatic nerve fascicles rotate along the rotation axis. **w.** same schemes as in A, C showing fibers vertical. **x.** registered views of the same fiber shown in **d**, now with a rather constant scattered signal apparently independent of the angle view. Bottom: color scale and intensity levels. **y.** Raw XZ image of a single plane, single view in the fascicle. **z.** The same image with angle/view color coding, showing scrambled colors and no specific pattern. **aa.** angle color coding legend of views a1 to a10. **ab,ac.** same as in R-V, raw grayscale YZ view and its angle color-coded version. Scale bars: **b**, 0.3 mm; **m**, 0.5 mm; **r**, 0.2 mm.

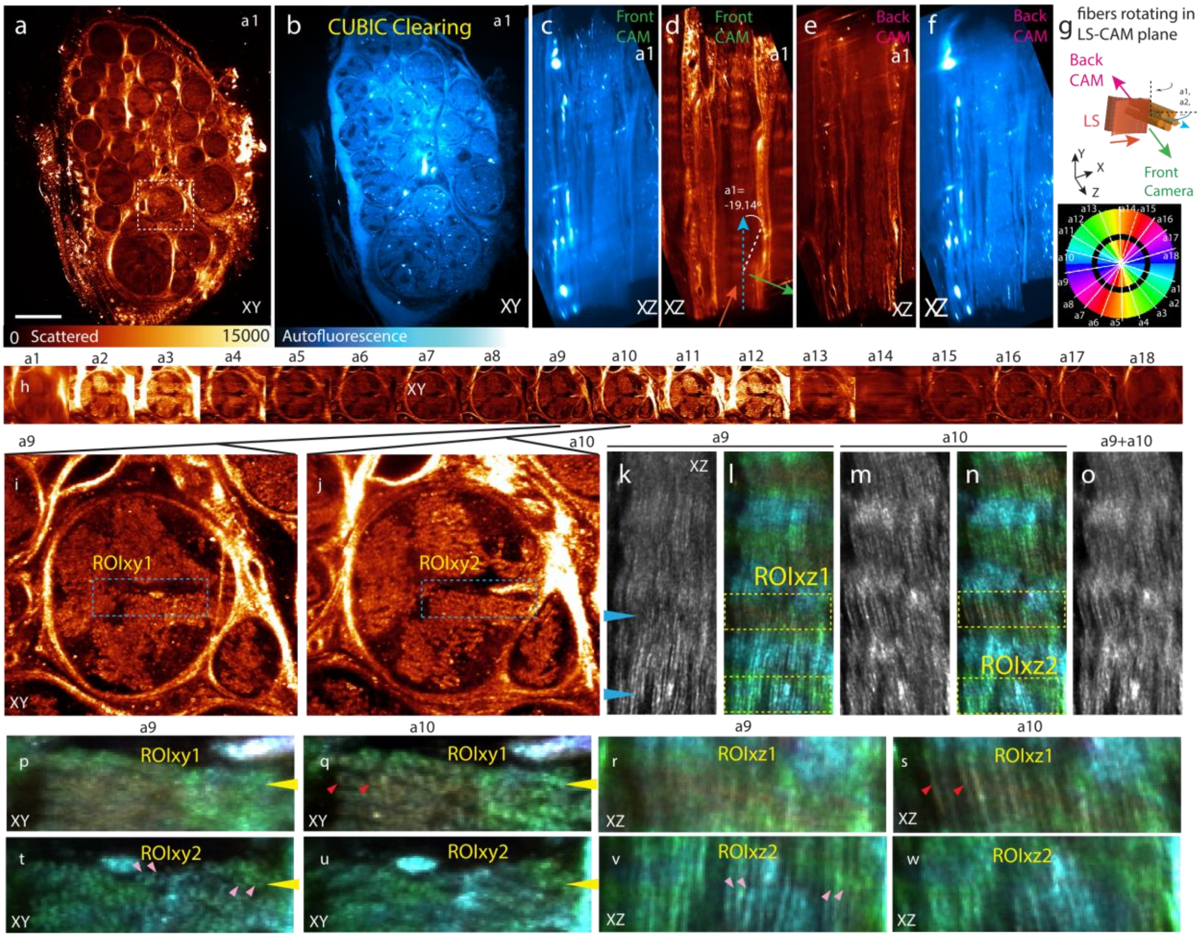


**Supplementary Figure 3**: **Orientation-dependent scattering in CUBIC-cleared sciatic nerve fibers. a,b.** XY images of LSSM signal and Autofluorescence with color intensity scale underneath. **c-f.** Transverse XZ sections (both channels, both cameras) along a fascicle shown on A with dashed square for a view where fascicle marks an angle a1 to the fiber axis. **g.** 3D scheme of light-sheet to fibers orientation: fibers rotate perpendicular to light-sheet-detection plane (similar to **Suppl. Fig. 2a-v**). Bottom: color coding scale for angular rotations of the 18 views a1-a18. **h.** Registered cross section XY of dashed square for all views showing again an LSSM signal increase and a half-turn redundancy (i.e. two high intensity peaks at about 180° difference). Noticeably, a strong LSSM signal appears in the perineurium that was not visible in the MeOH:BABB cleared sample (Suppl. Fig. 2**a-v**). **i,j:** a9 and a10 views show the best contrast on the fascicle bundle at the selected Z position, corresponding with a geometrical alignment of the fibers with the detection optical axis Z, whereas the other views are affected by the clearing deficiencies. **k,m:** Transverse XZ images of view a9 and a10, within the regions ROIxy1 and ROIxy2, respectively. **l,n:** color coding of all views applied to K and M respectively (see Supplementary Methods 2 for description). In a similar fashion to what shown in **Suppl. Fig. 2**, angular dependent LSSM signal forms a patchy pattern along the fiber axis. Here, however, individual fibers are discernable and seem to have different orientations according to the Y position. **o.** Sum of **k** and **m**, showing fiber buckling similar to **Suppl. Fig. 2. p-q, t-u:** angle color coding applied to ROIxy1 and ROIxy2. **r,s** and **v,w:** zoomed regions from ROIxz1 and ROIxz2 in L and N. Fibers in **s** correspond to rings in **q** (red arrowheads), and fibers in **v** correspond to rings in **t** (pink arrowheads). Rings resemble well-known myelinated axons in horse sciatic nerves.

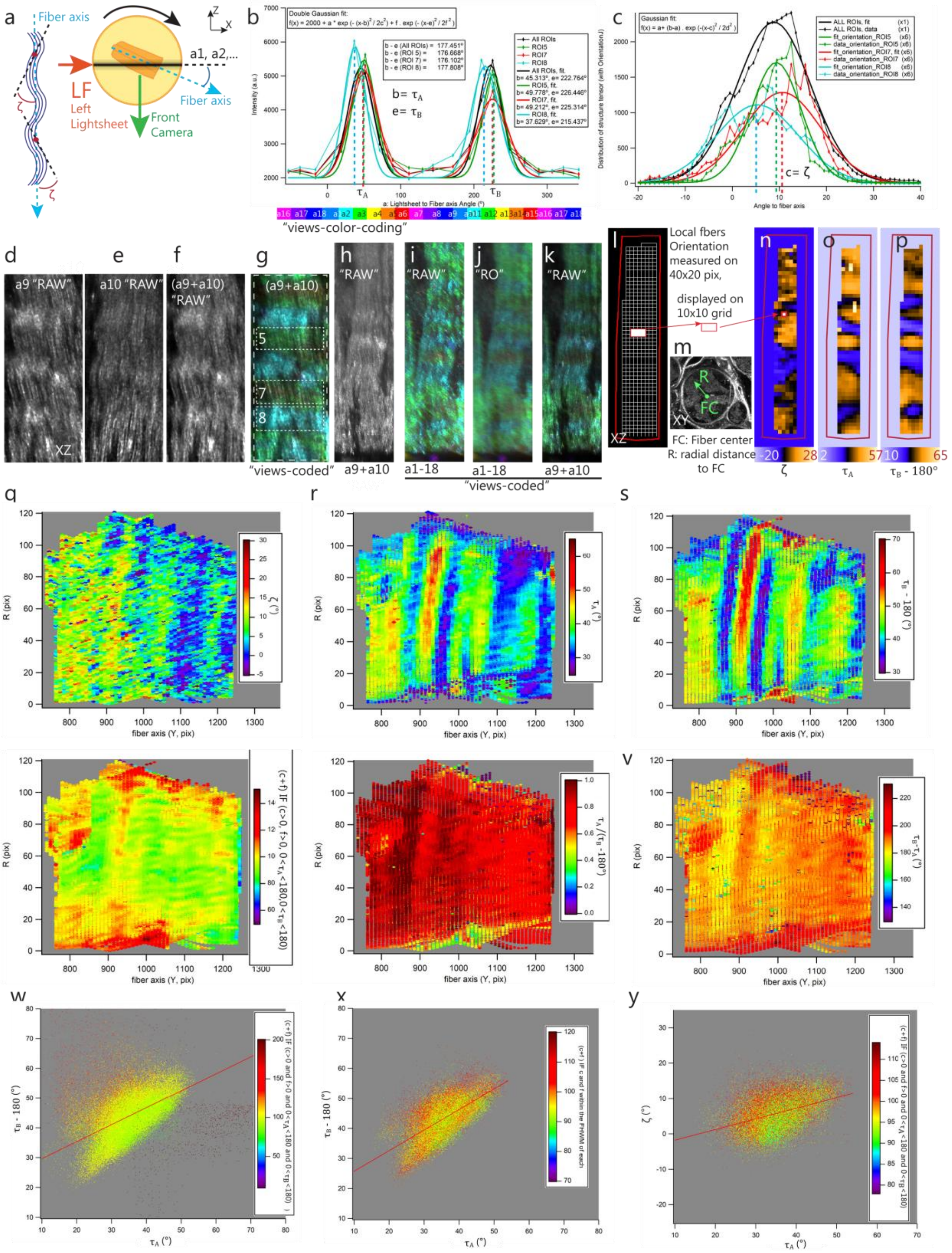


**Supplementary Figure 4**: **In CUBIC cleared sciatic nerve, local fascicle fibers orientation correlates with mvLSSM signal.** **a.** Left: scheme representing the observed buckling of nerve fascicle with an angle ζ describing the local tangent to the main fiber axis. Right: light-sheet (LS) orientation describing an angle a1, a2, …, a18 (18 views) with the fiber's main axis, sample rotation perpendicular to LS-detection plane, same sample and data as in **Suppl Fig. 3**. **b.** LSSM intensity measured in different regions of interest (ROI) from **g**, with color coding below the graph corresponding to colors in **g**, **i-k**. Two peaks are detected, approximately spaced 180° (see values in box). Double gaussian fitting (see equation in inset, parameters {a,b,c,d,e,f}) yields two angular values b= $\tau_A$ and e= $\tau_B$. Data and fit curves are colored with the dominant angular colors of ROI 5 (green), 7 (red), 8 (cyan) in **g**. Note how shifted is the cyan curve. Black data and fit curve stand for the intensity of the larger region including ROIs 5-7-8, i.e. the full image **g** (see large, dashed rectangle). **c.** Plot of the structure tensor, i.e. local orientation distribution, measured (see supplementary note 2) in the same ROIs on signal of image in **f**, being the sum of **d** (view a9) and **e** (view a10). Distributions are gaussian-fitted (see inset for equation) to yield a center angle ζ. **d-g**: a small XZ portion of fascicle shown in Supp. Fig 4. **h-k**: a larger portion of the same fascicle, from a different Y plane, where volume analysis is made (analysis images **l** to **y**). **h.** shows the raw intensity of views a9+a10, **i.** the color-coded projection of all views (noisy, with bright scatterers from specific views), **j.** Color coded projection with a previous low pass filter ("remove outliers" in Fiji) to better show color sections, and **k.** Color codes intensities of **h** with the colors distribution of **j**. The angles $\tau_A$, $\tau_B$ and ζ were measured by the above fitting strategies (in **b**,**c**) across the whole volume of the fiber region shown in **h-k**. **l**. Analysis layout: the volume is divided into a grid of 10 pixels pitch and measurements made on 40x20 regions for each grid position. **m.** An XY image of the fiber cross section shows the Fiber center (FC) and the vector R used for radial position in plots **q** to **v**. **n,o,p**: Heat maps of ζ, $\tau_A$ and ($\tau_B$ -180°) in selected Y plane, showing strong similarity between LSSM peaks (**o** and **p**) and milder similarity of both with ζ (see also color scales below images). **q** to **v**: Plots report different values in color heatmaps for the whole fiber volume, i.e. bottom axis is fiber axis Y and left axis is the radial position **r** (drawn in **m**) inside the fiber. **q:** angle ζ. **r:** Angle $\tau_A$. **s:** Angle $\tau_B$ -180°. **t:** sum of {c+f} parameters from **b** expressing sharpness of the LSSM angular signal with the sum of the gaussian widths (green: sharper peaks sum, red: wider peaks sum). Data points of {c+f} were filtered for physical meaningfulness to discard aberrant fits (see Supplementary Note 2). **u**: ratio of $\tau_A$ over [$\tau_B$ -180°] generally close to 1, V: difference $\tau_B$ - $\tau_A$ generally close to 180°. **w.** plot of all data points filtered similarly to **t**, of [$\tau_B$ -180°] vs. $\tau_A$, with color coding showing peaks sharpness (c+f). **x.** The graph shows the same for values of {c} and {f} within the full-width-at-half-maximum (FWHM) of their distribution, hence discarding outliers and representing only sharp scattering peaks. The linear relation between $\tau_A$ and $\tau_B$ is clear, showing the two LSSM peaks move together. **y**. plots ζ vs. $\tau_A$, showing a milder linearity, hence suggesting a mild relation, within the limits of the data spread, between local buckling and scattering orientation.

### Supplementary Methods 2. Sciatic nerve image processing

#### Registration of sciatic nerve data

Multiview mvLSSM volumes registration of sciatic nerve data was performed with plugins available in Fiji[7] with the aim to register only a subvolume (centered on one fascicle) for multiview intensity quantification along a specific part of fiber. In brief, considering multiview XYZ volumes with Y the sample's axis of rotation, the volume was cropped in Y to conserve only the fiber of interest, and turned with TransformJ (around X, 90°). The main fiber axis angle was measured manually for each view and each volume view rotated accordingly to obtain all views with the fiber of interest parallel. For each XZ plane, "rigid body" registration with StackReg was performed along the 18 angle views to align all to the first angle view. This provided a coarse alignment that was then refined with a second "rigid body" run of StackReg, this time along the 3rd dimension (in Y for the original volume). The same procedure was applied to both datasets (BABB-cleared and CUBIC cleared). This procedure enabled satisfactory alignment of images under the assumptions that i} quick registration was the priority to circumvent memory issues (because of cropped volumes) and ii} assumed that the Y rotation axis of the sample is well aligned with the Y axis of the volumes.

#### Image processing for visualization of angular views in sciatic nerves

In Supplementary Fig 2, 3 and 4, color rotation encoding was performed by "temporal color-code" plugin in Fiji, with "BIOP12colors.lut" after swapping z to t frames. Supplementary Fig. 2r, 2z, 2ac and 4j used raw LSSM data which exhibit high intensity foci. To smoothen images for better color perception, in Supplementary Fig. 2m-p, 2s-t and 4k, images were low-pass filtered with "remove outliers" (both for high and low intensity pixels, i.e. "dark" and "light" background) with radius 2 pixels, intensity 50. However, for the other images (Supplementary Fig. 3l, 3n, 3p-w, 4g and 4l) where the goal was to represent the best view where internal fibers are best resolved (e.g. panels Supplementary Fig. 3k, 3m, 4f 4i) with rotation-encoded colors, the color projection of raw data would degrade the contrast due the fact that some of the views were blurred by the partial clearing (i.e. by rotating the sample, the optical path increases through the sample, hence the same location is not imaged with the same clarity among all views if clearing is partial). To circumvent this, those images representing the best view that show internal fibers structures were colored with the aforementioned low-pass filtered image, let call it {RGB}. To do so, RGB is split, each component turned to 32-bit and multiplied by the original image of interest, then recombined into a new RGB image.

### Supplementary Methods 3. Modeling framework: Light scattering by infinite cylinders

#### Light scattering by neural fibers

Neural fibers longer than their diameter can be modeled by infinite circular cylinders. For such a model, the interaction and scattering of a plane electromagnetic wave with a wavelength shorter than the cylinder length is well described[8,9]. Unlike spherical scattering (Mie theory), where scattering occurs in all directions, an infinite cylinder scatters light into a cone. Thus, an intensity is only observed by a detector if it "looks" in the direction where the scattered light is propagated.

To study the scattering problem, we first must define the coordinate system. In the microscope coordinate system (*XYZ*), the light-sheet propagates along the X-axis, $\boldsymbol{r}_i = (1,0,0)$, until it is incident on a cylinder (fiber) with orientation $\boldsymbol{r}_f$. Additionally, the detector is placed at the Z-axis and modeled as a unit vector $\boldsymbol{r}_o = (0,0,1)$ (Supplementary Fig. 5b).

When the propagating light $\boldsymbol{r}_i$ is incident on the fiber $\boldsymbol{r}_f$, the scattered light spreads along the surface of a cone, whose apex is the point of incidence, and the cone half-angle is the angle between $\boldsymbol{r}_i$ and $\boldsymbol{r}_f$. This angle is $\frac{\pi}{2} - \varphi$, where φ is named the elevation angle; thus, φ represents a cone with a specific aperture. Additionally, we should consider that the scattered light waves are not equal for the different propagation directions in the cone. Thus, we identify the angle around the cylinder's cone through the scattering angle, θ (Supplementary Fig. 5a). To mathematically characterize θ, we define the plane containing $\boldsymbol{r}_i$ and $\boldsymbol{r}_f$ as the plane of incidence. Then, we denote the scattered direction of interest within the cone as $\boldsymbol{r}_s$. The plane containing $\boldsymbol{r}_s$ and $\boldsymbol{r}_f$ forms the scattering plane. The scattering angle, θ, is the angle between the plane of incidence and the scattering plane. For θ = 0°, the scattering plane contains both $\boldsymbol{r}_i$ and $\boldsymbol{r}_f$.

The cone-shaped scattering propagation implies that the model only predicts a measurable signal intensity when the cone of scattered light is tangential to the detector normal, such that a scattered wavefront $\boldsymbol{r}_s$ is directed toward the detector ($\boldsymbol{r}_s = \boldsymbol{r}_o$). Consequently, this only happens when both the detector normal $\boldsymbol{r}_o$ and the direction of the incident light $\boldsymbol{r}_i$ form the same angle with the fiber direction $\boldsymbol{r}_f$ (Supplementary Video 4). To solve the forward problem, we must identify the relationship between the fiber orientation $\boldsymbol{r}_f$, the scattered cone aperture associated with the elevation angle φ, and the specific propagation direction of the cone scattered towards the detector identified by θ. The following subsection derives the relationship between $\boldsymbol{r}_f$, φ, θ to simulate the scattering profiles. Later, we outline the standard formalism (based on the text from Bohren and Huffman) to characterize the relationship between the incident and the scattered electric field vector in the scattering process, which enables us to estimate numerically the scattered intensities. The last subsection exemplifies some consequences of the derived equations in the macroSPIM system (Y-axis sample rotation), including light polarization considerations.

#### Geometrical derivations in the microscope space

The fiber direction $\boldsymbol{r}_f$ can be described in spherical coordinates by the two angles $\alpha_f$ (tilt of the fiber with respect to the rotation axis) and $\beta_f$ (azimuthal angle in the perpendicular plane). The relationship between the fiber direction and φ is derived from the definition of the elevation angle:

$$\boldsymbol{r}_f \cdot \boldsymbol{r}_i = \cos(\pi/2 - \varphi) = \sin(\varphi) \quad \text{[Eq. 1]}$$

As exposed earlier, for the scattering to be detected, the scattered wavefront $\boldsymbol{r}_s$ should be directed towards the detector, $\boldsymbol{r}_s = \boldsymbol{r}_o$. Under these conditions, using the dot product, we have:

$$\boldsymbol{r}_f \cdot \boldsymbol{r}_i = \boldsymbol{r}_f \cdot \boldsymbol{r}_o$$

To evaluate the scattered amplitude, we apply the definition of the elevation angle:

$$\boldsymbol{r}_f \cdot \boldsymbol{r}_i = \boldsymbol{r}_f \cdot \boldsymbol{r}_o = \cos(\pi/2 - \varphi) = \sin(\varphi)$$

And due to the detector and light-sheet being orthogonal, $\boldsymbol{r}_o \cdot \boldsymbol{r}_i = 0$. Consequently, θ has a simple relation to φ, which we determine using the definition of θ as the angle between the plane of incidence and the scattering plane:

$$\cos(\theta) = \frac{(\boldsymbol{r}_i \times \boldsymbol{r}_f)\cdot(\boldsymbol{r}_o \times \boldsymbol{r}_f)}{|\boldsymbol{r}_i \times \boldsymbol{r}_f||\boldsymbol{r}_o \times \boldsymbol{r}_f|} = -\frac{\cos^2(\pi/2-\varphi)}{\cos^2(\varphi)} = -\tan^2(\varphi) \quad \text{[Eq. 2]}$$

where × represents the cross-product. Once φ and θ are known, the expected scattering amplitude is determined by the standard formalism, summarized in the next subsection.

#### Scattered electric field and intensity derivations

The scattered electric field is calculated using the exact solution for an infinite cylinder. The scattered electric field ($E_s$) is related to the incident field ($E_i$) via the 2×2 Amplitude Scattering Matrix ($T$):

$$\begin{pmatrix} E_{\parallel s} \\ E_{\perp s} \end{pmatrix} = f(\varphi, k, r, z) \begin{pmatrix} T_1 & T_4 \\ T_3 & T_2 \end{pmatrix} \begin{pmatrix} E_{\parallel i} \\ E_{\perp i} \end{pmatrix} \quad \text{[Eq. 3]}$$

where the propagation factor $f(\varphi, k, r, z)$ is defined as:

$$f(\varphi,k,r,z)=e^{i3\pi/4}\sqrt{\frac{2}{\pi kr\cos\ \varphi}}e^{ik(r\cos\ \varphi+z\sin\ \varphi)}$$

The elements $T_1, T_2, T_3$, and $T_4$ form the amplitude scattering matrix, which depends on θ and φ while accounting for the fiber's refractive index and radius. $E_{\parallel}$ and $E_{\perp}$ represent the parallel and perpendicular components of the electric field with respect to the plane of incidence. The decay factor includes the factor $\sqrt{2/(\pi kr\cos\ \varphi)}$ to correct for the geometric decay of wave intensity as a function of the perpendicular distance from the fiber axis ($r$) and the cone aperture angle (φ). The complex exponential terms indicate propagation phase and do not influence intensity measurements.

Each T-matrix element describes field amplitudes as complex numbers and can be estimated numerically[8–10]. Supplementary Fig. 5c shows absolute values of each scattering element for all possible θ, φ, and the green dashed line indicates which values can be observed by fibers in the system with Y-sample-rotation. In other words, each value in the green line corresponds to a specific fiber orientation, (α, β), where the scatter is measurable, and its orientation is related to a specific (θ, φ) through Equations 1 and 2.

To correlate the complex field amplitudes with experimental data, we need to estimate the total scattered intensity. This requires decomposing the incident wave $E_i$ into components relative to the plane of incidence (the plane containing $\boldsymbol{r}_f$ and $\boldsymbol{r}_i$):

*Case 1: Linearly Polarized Light*

For linearly polarized light, the rotation axis ($Y$ in the macroSPIM system) serves as the reference. The direction of polarization forms an angle ψ with the rotation axis, where ψ = 0° corresponds to s-polarization and ψ = 90° to p-polarization, consistently with Supplementary Fig. 1n. The incident polarized vector in the laboratory reference frame is:

$$\boldsymbol{E}_i^{lab}=(0,\cos\psi,\sin\psi)$$

To decompose this field into local cylinder basis vectors, the perpendicular basis vector ($\hat{e}_{\perp}$) to the incidence plane is obtained via the cross product:

$$\mathbf{N}=\boldsymbol{r}_f\times\boldsymbol{r}_i\ \text{ and }\ \hat{e}_{\perp}=\frac{\mathbf{N}}{|\mathbf{N}|}$$

The parallel vector ($\hat{e}_{\parallel}$) is then:

$$\hat{e}_{\parallel} = \hat{e}_{\perp} \times \boldsymbol{r}_i$$

Projecting $\boldsymbol{E}_i^{lab}$ onto these vectors yields the incident components $E_{\parallel}^{i} = \boldsymbol{E}_i^{lab} \cdot \hat{e}_{\parallel}$ and $E_{\perp}^{i} = \boldsymbol{E}_i^{lab} \cdot \hat{e}_{\perp}$. The total intensity is proportional to the square of the scattered field (applying Equation 3):

$$I_{total} \propto |\vec{E}_{total}|^2 = |E_{\parallel}^{s}|^2 + |E_{\perp}^{s}|^2 \text{ [Eq. 4]}$$

*Case 2: Unpolarized Light*

Unpolarized light is treated as a 50/50 incoherent mix of parallel and perpendicular incident light. The total scattered intensity is the average of these contributions:

$$I_{\text{unpol}} \propto \frac{1}{2} \cdot \left(\frac{2}{\pi \,\text{krcos}\,\varphi}\right) \cdot [(|T_{11}|^2+|T_{21}|^2)+(|T_{12}|^2+|T_{22}|^2)] \text{ [Eq. 5]}$$

Supplementary Fig. 5d,e exemplifies the application of Equation 5 for all (Supplementary Fig. 5d) and selected (Supplementary Fig. 5e) θ and φ pairs.

#### Derivation of Scattering Profile Features

Once the relationship between the fiber orientation (α, β), the scattering cone geometry (θ, φ) and the scattering intensities is known. We can simulate the scattering profiles. In this subsection, we show derivations for the macroSPIM system ($Y$-axis sample rotation). Further insights into the Z-axis sample rotation (Ultramicroscope system) are provided in Supplementary Note 3.

In the macroSPIM system, the fiber orientation $\boldsymbol{r}_f$ is represented as:

$$\boldsymbol{r}_f = [\sin\alpha_f \cos\beta_f \,, \cos\alpha_f \,, \sin\alpha_f \sin\beta_f]$$

For detection where $\boldsymbol{r}_i = [1,0,0]$ and $\boldsymbol{r}_o = [0,0,1]$, the condition of detection, $\boldsymbol{r}_f \cdot \boldsymbol{r}_i = \boldsymbol{r}_f \cdot \boldsymbol{r}_o$ leads to:

$$\sin\alpha_f \;\cos\beta_f \; = \sin\alpha_f \sin\beta_f \,; \tan\beta_f = 1; \;\beta_f = \pi/4 + n\pi, n \in \mathbb{N}$$

This geometric derivation supports the observed 180° periodicity in experimental results, as well as the 90° difference between the applied rotations for measuring a peak for different light-sheets/detector positions (i.e., cases $\boldsymbol{r}_i = [-1,0,0]$ and/or $\boldsymbol{r}_o = [0,0,-1]$).

Furthermore, incorporating scattered intensity allows for the correlation of peak intensity with fiber tilting ($\alpha$), as well as the polarization of the incident light-sheet (Equation 4 for linearly polarized light or

Equation 5 for unpolarized light). Predictions indicate that in-plane fibers ($\alpha = 0°$) yield larger scattered intensities than out-of-plane fibers ($\alpha = 90°$) regardless of polarization (Supplementary Fig. 5f-h). Simulations confirm that s-polarization ($\psi = 0°$) maximizes scattered intensity, while p-polarization ($\psi = 90°$) significantly reduces it (Supplementary Fig. 5g), in agreement with the experimental observations (Supplementary Fig. 1o).

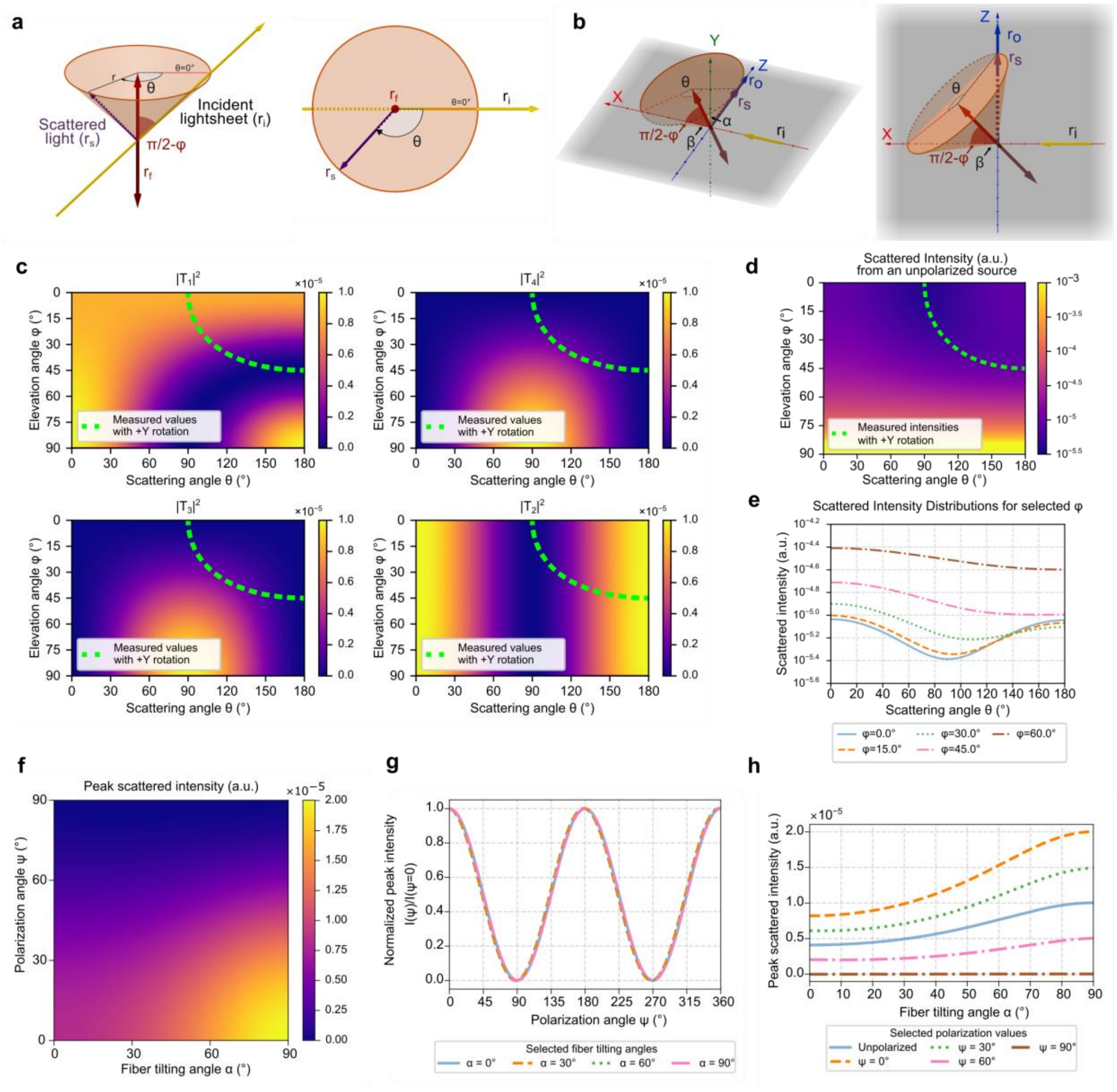


**Supplementary Figure 5**. **Theoretical framework of scattering by infinite cylinders and its relationship with the scattered intensities.** a. Geometry of light scattering by an infinite cylinder. The scattered light forms a conical sheet coaxial with the cylinder axis. The half-angle of the cone is the angle between the cylinder axis and the incident ray. Definitions of the elevation (φ) and scattering angles (θ) are given in text. b. Geometry of the light scattering within the microscope elements context. c. Magnitude values of the scattering amplitude matrix elements for the different θ and φ angles. The green dashed lines indicate which values can be measured in the microscope with Y-sample rotation. d,e. Distribution of the scattered intensities from an unpolarized source for every possible cone of scattered light (e), and for selected cone sheets (f). f. Dependence of the peak-scattered intensity detected by the macroSPIM system on fiber tilting (α) and incident light polarization (ψ). g. Variation of the peak scattering intensity relative to s-polarization (ψ=0°) when changing the linear polarization for different fiber tilting. h. Peak-scattered intensities for selected polarization angles and unpolarized light (ψ) depending on fiber tilting (α).

### Supplementary Note 3. Scattering imaging and modeling through Z-axis rotation in an Upright "Ultramicroscope".

The choice of rotation axis for mvLSSM acquisition is often constrained by gravity, making a vertical axis the most practical option. To evaluate this configuration, we acquired mvLSSM data using an Ultramicroscope II system (Miltenyi/LaVision BioTec GmbH, Germany), which features upright detection and left-side illumination (Extended Data Fig. 4a). In this setup, the rotation axis aligns with the detection axis (Z-axis), producing scattering profiles distinct from those obtained with macroSPIM.

#### System and acquisition specifications

The system is equipped with a standard incoherent supercontinuum white laser (SuperK Extreme, NTK Photonics, Germany). The optical configuration includes dual-sidesheet illumination, seven-slot filter wheels, an Olympus MVX10 zoom microscope body (Olympus, Japan) with a 2×/NA 0.5 Olympus MVPLAPO objective lens (1.26×–12.6× magnification with 10-fold internal zoom and 0.63x tube lens) fitted with a custom dipping cap (LaVision BioTec GmbH). Image detection was performed using a 5.5 MP Andor Neo sCMOS camera (Oxford Instruments GmbH, Germany), featuring a sensor size of 16.6 × 14 mm (2560 × 2160 pixels).

Brain samples were imaged at 1.26× magnification, and voxel size 2.6×2.6×5.16 $\mu m^3$. DBE was used as the imaging medium. Whole-brain volumes were acquired across 16 views by rotating the sample through [0°, 360°) using a custom holder, with 620 nm illumination and top detection.

#### Scattering profiles features

For mvLSSM Z-axis-rotation data, simulations indicated that peak position depends on fiber azimuth (βf), while peak amplitude and spacing between peaks depend on tilt ($\alpha_f$) (Extended Data Fig. 4b). For fibers perpendicular to the axis of rotation $\alpha_f = 90°$, the peaks are separated by 180°, but this distance decreases until they merge for $\alpha_f = 45°$. Below this threshold, amplitude diminishes toward a flat profile (Extended Data Fig. 4c,e–g).

We found the expected peak patterns in experimental data (Extended Data Fig. 4d,h-k). We observed that regions of interest in the corpus callosum (Extended Data Fig. 4h) and the fimbria (Extended Data Fig. 4i) showed varying distance between the peaks, suggesting different tilting of the tracts. Similarly, in cingulum, or in caudoputamen bundles, we observed a single peak, suggesting a tilt $\alpha_f > 45°$ (Extended Data Fig. 4j-k).

These observations imply that azimuth and partial tilt ($\alpha_f \in [45°, 90°]$) can be estimated from peak positions in a single mounting position. However, the lack of periodicity requires full [0°, 360°) coverage and precludes PCA-based peak detection, reducing azimuthal resolution and complicating peak-finding approaches. Alternative methods typically depend on hyperparameter tuning and may be more susceptible to noise. Additionally, the lack of periodicity complicates the possibility of solving crossing fibers, since the identification of peaks from the same fiber bundle and differentiating them from single-peak fibers will likely not be trivial in experimental data.

#### Theoretical relationship between the scattering profile features and the fiber orientation.

To determine the relationship between the scattering signal profile and in-plane orientation, we begin with the peak measurement condition. Let $\boldsymbol{r}$ be the fiber direction at the peak position and $\boldsymbol{r}_f$ be the original fiber direction, such that:

$$\boldsymbol{r} = R_z(\omega_{\text{peak}})\boldsymbol{r}_f$$

where $R_z$ represents the right-hand rotation around the Z-axis and $\omega_{\text{peak}}$ is the rotation angle. The peak condition is defined by:

$$\boldsymbol{r}_i \cdot \boldsymbol{r} = \boldsymbol{r}_o \cdot \boldsymbol{r}$$

In spherical coordinates, the fiber direction is expressed as: $\boldsymbol{r} = [\sin(\alpha)\cos(\beta), \sin(\alpha)\sin(\beta), \cos(\alpha)]$

Solving for the angular relationship:

$$\cos(\beta) = \frac{\cos(\alpha)}{\sin(\alpha)} = \frac{1}{\tan(\alpha)}$$

$$\beta = \arccos\left(\frac{1}{\tan(\alpha)}\right)$$

This expression implies that: (i) no solution exists for $\alpha > 135°$or $\alpha < 45°$; (ii) the mid-point between the peaks indicates the rotation needed to align the fiber with the incident light ($\omega_{\text{midpoint}}$).

We can leverage the insight (ii) to recover the in-plane orientation. Such that

$$\boldsymbol{r_i} = R_z(\omega_{\text{midpoint}})\boldsymbol{r}_f{}^{\text{in plane}}$$

Since we know the in-plane position, and the necessary sample rotation to measure a peak ($\omega_{\text{peak}}$), we can use:

$$\beta = \beta_f + \omega_{peak}$$

Such that β represents the specific in-plane orientation of the fiber when it is yielding a peak intensity. Thus, the tilting is:

$$\alpha = \arctan\left(\frac{1}{\cos(\beta)}\right)$$

Once known $\alpha$ and β, we reconstruct the original direction as:

$$\boldsymbol{r}_f = R_z(\omega_{peak})^T \boldsymbol{r}$$

This approach restricts fiber tilt reconstruction to the range $\alpha \in [45°, 90°]$. Confounding factors affecting amplitude modulation remain a common challenge across acquisition systems, meaning that full 3D reconstruction requires a second mounting to resolve these ambiguities. Nevertheless, partial tilt estimates from each mounting can help reduce uncertainties inherent to triangulation.

Despite these limitations, results obtained with the ultramicroscope confirm the robustness of orientation-dependent scattering as an intrinsic tissue property and demonstrate that the tLSSM framework can be adapted to different light-sheet microscopy platforms.

### Supplementary Methods 4. Registration pipeline

To model the scattering profile on a voxel-wise basis, we implemented a multi-resolution image registration pipeline using the open-source software Elastix version 5.0.1[11]. The pipeline was designed to align volumetric images acquired across different rotation angles and mounting positions.

**Preprocessing.** All volumetric images were downsampled by a factor of four in each spatial dimension to reduce computational load. To mitigate the influence of high-intensity scattering regions, a gamma correction with $\gamma = 0.5$ was applied to the intensity values prior to registration.

**Registration Strategy.** Unless otherwise stated, affine transformations were estimated by maximizing the mutual information between fixed and moving images. Mutual information was computed using 32 histogram bins. The registration was performed at three resolution levels using a smoothing image pyramid with downsampling factors of 4, 2, and 1, respectively. At each resolution level, the adaptive stochastic gradient descent optimizer was used for 1000 iterations, with 65536 new samples drawn per iteration.

**Intra-Series Alignment.** Each volume $I_n$ in the rotation series (where $n \geq 1$) was registered to the previously acquired volume $I_{n-1}$, estimating a transformation $t_n(x): I_n(x) \rightarrow I_{n-1}(x)$. The resulting transformations were composed recursively to register each volume to the first volume in the series, $I_0$, which was acquired using the back camera and left light-sheet at the 0-degree mounting position. This volume served as the *reference volume* for all subsequent steps.

**Inter-Series Alignment.** The first volume of the 90-degree rotation series was registered directly to the reference volume. The resulting transformation parameters were then applied to all subsequent volumes in the 90-degree series, ensuring consistent alignment across datasets acquired at different mounting positions.

**Final Refinement.** After concatenating all estimated transformations, they were applied to the original (non-downsampled) voxel-size volumes. A final refinement step was performed by registering each volume to the reference volume at full resolution and original intensity range. This step used 2000 iterations of the ASGD optimizer without an image pyramid, allowing for fine-scale alignment.

### Supplementary Methods 5. Sample preparation

#### Mice

The animals were housed in groups of four in a 12-hour light/dark cycle with ad libitum access to a standard diet until the day of surgery. Following the surgical intervention, the animals were housed individually, maintaining the same conditions of light and diet. In all cases, the temperature was kept constant at 22°C, while the relative humidity ranged between 30% and 80%, although it was not strictly controlled.

#### Induction of localized demyelination in the corpus callosum

To induce demyelination in the corpus callosum of wild-type mice, we performed intracranial injections of 1 μl of 1% lysophosphatidylcholine (LPC, Cat. No. L4129, Sigma-Aldrich, St. Louis, MO, USA) in sterile saline. LPC is a bioactive lipid derived from phosphatidylcholine that targets oligodendrocytes, disrupting their cellular membranes and leading to their subsequent death. This initial loss of oligodendrocytes triggers a localized inflammatory response, resulting in controlled focal demyelination.

#### Surgical procedure and postoperative care

The animals were anesthetized with isoflurane delivered through a calibrated vaporizer (induction at 4–5%, maintenance at 1–2%). The depth of anesthesia was confirmed by the absence of reflexes, including the corneal and withdrawal reflexes. Once adequate anesthesia was ensured, the animals were secured in a stereotaxic frame. Throughout the procedure, vital parameters, including body temperature, heart rate, and oxygen saturation, were monitored to ensure physiological stability. The surgical site was disinfected thoroughly before administering local anesthesia via subcutaneous injection of lidocaine, along with preoperative analgesia using buprenorphine. Ophthalmic gel was applied to protect the eyes from desiccation and damage. A sagittal incision was made in the scalp to expose the skull. The coordinates for the corpus callosum (AP: 0.97; DV: 2.6; ML: 1.00) were identified using a stereotaxic atlas and marked on the skull. For the H&E lesion study (AP: 0.85; DV: 2.35; ML: 0.50) coordinates were used. Burr holes were drilled at the designated coordinates using a high-precision stereotaxic drill. After drilling, a microinjection needle was carefully inserted into the target region, and 1 μl of 1% LPC was injected at a controlled rate of 0.1 μl/min to minimize mechanical damage to surrounding tissues. Following the injection, the needle was left in place for 10 minutes before being slowly withdrawn over an additional 10-minute period to reduce tissue disruption. The incision was closed with absorbable sutures, and the animals were transferred to a temperature-controlled recovery chamber. Post-surgical monitoring continued until the animals fully regained consciousness. During the first three days post-surgery, the animals were monitored twice daily to ensure wound integrity and to prevent infections. Additional analgesia was administered with buprenorphine, and antibiotics were provided to prevent infections.

#### Lesion assessment and tissue collection

The effectiveness of demyelination was evaluated using magnetic resonance imaging (MRI) performed 7 days after injection. This analysis confirmed that the lesion was localized to the corpus callosum and exhibited consistent size across all animals, meeting experimental objectives. Once the desired lesion size was verified, the animals were sacrificed via an overdose of pentobarbital and transcardially perfused with 4% paraformaldehyde (PFA) or glioxal in heparinized saline. After perfusion, the brains were carefully extracted and stored appropriately for subsequent analysis.

The mouse used for H&E staining underwent similar procedures with the following exceptions: terminal anesthesia prior to perfusion was achieved using a Hypnorm–midazolam mixture; and 10% neutral buffered formalin was used as fixative.

#### Horse sciatic nerve

The tissue samples used in this study were taken from a 15-years-old Danish warmblood horse. The horse was euthanized by a captive bolt pistol followed by exsanguination. After euthanasia the sciatic nerve was carefully dissected from the gluteal region of the left hindlimb and followed for approx. 30 cm down the back of the thigh and cut before splitting into the N. tibialis and N. peroneus communis. The sample was embedded into 4% PFA and stored at 4°C until further analysis.

#### Haematoxylin and eosin (H&E) staining

An LPC-injected mouse brain was embedded in paraffin and sectioned coronally at 10 µm using a manual microtome (RM2125 RTS, Leica Biosystems, Germany). Tissue sections were mounted onto glass slides (Starfrost, Hounisen, Denmark) and dried in the oven for 20 mins at 60 °C. Paraffin was removed using Histo-Clear (two 5-minute changes). The sections were then rehydrated by sequential immersion of the slides in 100% ethanol (two 2.5-minute changes), followed by 96% and 70% ethanol (one 2-minute change each), and subsequently rinsed in running deionised water (5 mins). H&E staining was achieved as follows: immersion of the slides in haematoxylin (4 mins), rinsing in running deionised water (10 mins), and immersion in eosin (1 min). Afterwards, the sections were dehydrated with one dip of the slides in 70% alcohol, immersion in 96% alcohol (2 mins) and 100% alcohol (two 2.5-minute changes), and then cleared in xylene (two 2.5-minute changes). Finally, the sections were cover-slipped with Tissue-Tek® Glas Pertex (Sakura, Japan), scanned at 20x by a VS200 slide scanner (Olympus, Japan), and imaged with the Olympus Viewer plugin (v2.4.1) for Fiji (Imagej2 v2.16.0/1.54p)[7].

#### iDISCO+: Tissue processing and immunolabeling

##### Tissue pretreatment

Residual fixative (PFA or glyoxal) was removed by consecutive washes in 1X phosphate-buffered saline (PBS, 10X, Ambion, Cat. No. AM9624, Thermo Fisher Scientific, Waltham, MA, USA) for 30 minutes. Samples were progressively dehydrated using a methanol series (Prod. No. 1.06009, Merck-Sulpeco, Germany) in water, with graded concentrations of 20%, 40%, 60%, 80%, and 100% methanol. Each step consisted of 60-minute incubations at room temperature. Once dehydration was completed, the tissues underwent delipidation overnight at room temperature in a solution of 66% dichloromethane (DCM, Prod. No. VWRC83682.290, Avantor-VWR Chemicals, PA, USA) and 33% methanol under constant agitation. Following delipidation, tissues were washed twice in 100% methanol at room temperature and chilled at 4°C. They were then incubated overnight at 4°C in freshly prepared 5% hydrogen peroxide (H2O2) in methanol (Sigma, Cat. No. 216763-100ML, Merck KGaA, Darmstadt, Germany) to reduce residual pigmentation and enhance tissue clarity. After bleaching, tissues were rehydrated through a descending methanol series (80%, 60%, 40%, 20%) and finally in PBS, with 1-hour incubations per step at room temperature. Pretreatment concluded with two consecutive washes in PBS supplemented with 0.2% Triton X-100 (Sigma, Cat. No. X100-500ML, Merck KGaA, Darmstadt, Germany) for 1 hour at room temperature.

##### Immunolabeling and clearing

To facilitate antibody penetration, pretreated tissues were incubated for 2 days at 37 °C with gentle agitation in a permeabilization solution containing PBS, 0.2% Triton X-100, glycine (Sigma, Cat. No. G7126-500G, Merck KGaA, Darmstadt, Germany), and dimethyl sulfoxide (DMSO, Fisher, Cat. No. D128-4, Thermo Fisher Scientific, Waltham, MA, USA). This step was followed by a 2-day incubation at room temperature in a blocking solution containing PBS, 0.2% Triton X-100, 6% DMSO, 10% donkey serum (Jackson ImmunoResearch, Cat. No. 017-000-121, West Grove, PA, USA), and 1% bovine serum albumin (BSA, Sigma, Cat. No. A9647, Merck KGaA, Darmstadt, Germany) to minimize non-specific binding. Immunolabeling was performed by incubating the tissues with a fluorophoreconjugated primary antibody, mouse anti-MBP (BioLegend, Cat. No. 808408, San Diego, CA, USA), at a 1:500 dilution for 7 days at 37 °C under continuous agitation. After immunolabeling, samples were extensively washed in PBS containing 0.2% Triton X-100 to remove any non-specific antibodies. The clearing process was finalized by subjecting the tissues to a methanol series (20%, 40%, 60%, 80%, and 100%) at room temperature for 1 hour each step. Then, tissues underwent delipidation at room temperature for 3 hours in 66% DCM and 33% methanol, followed by two consecutive washes in 100% dichloromethane (Prod. No. VWRC83682.290, Avantor-VWR Chemicals, PA, US). Finally, tissues were incubated in dibenzyl ether (DBE, Sigma, Cat. No. 108014-1KG, Merck KGaA, Darmstadt, Germany), ensuring optimal tissue transparency and uniform clearing and used as final imaging medium.

**BABB vs. CUBIC clearing of horse sciatic nerve**

Two sections of the same sciatic nerve sample were used for comparative clearing with solvent-based MeOH:BABB (e.g. modified Murray's clear, Supplementary Fig. 2) and water-based CUBIC clearing (Supplementary Fig. 3,4). Imaging was performed between 1 week and 2 months after preparation.

With MeOH:BABB, a 4-5 mm dissected sample chunk was first embedded into a 1cm-diameter 2cm-long cylindrical agarose block, then dehydrated in steps of increasing grades of methanol (30 to 100%) during 24h and left at MeOH 100% for 48h. Samples were then immersed in BABB (Benzyl Alcohol-Benzyl Benzoate 1:2, Sigma Aldrich, 2412-1l and W213802-1KG-W) to match the refractive index until becoming transparent.

Another dissected nerve chunk was cleared with an updated CUBIC protocol[12]: the sample was directly immersed in CUBIC-L (a mixture of 10 wt% N-Butyldiethanolamine, Tokyo Chemical Industry, B0725, and 10 wt% Triton X-100, Merck 1.08603.1000), then incubated with shaking at 37 °C for 7 days (duration chosen based on the sample size). After 4 days, CUBIC-L was refreshed once. Following delipidation, the nerve was washed in PBS with gentle shaking at room temperature overnight. After that, it was embedded in an 1cm-diameter 2cm-long cylindrical agarose block to allow better manipulation during imaging, then immersed in 1:1 water-diluted CUBIC-R+ (a mixture of 45 wt% antipyrine, Tokyo Chemical Industry, D1876, and 30 wt% N-methylnicotinamide, Tokyo Chemical Industry, M0374) with gentle shaking at room temperature for 6 hours. The sample was then immersed in CUBIC-R+ at room temperature for 1-2 days before being ready for imaging.

### Supplementary Video 1

**Scattered signal by mvLSSM, axial travel through the mouse brain.** The video starts with maximum intensity projection of the autofluorescence signal (average of four views: front, back, left and right illuminations) to position the brain in axial orientation. In a specific plane half way (0:05), the 18 angular views are shown sequentially, followed (0:10) by the fractional anisotropic calculated from the latter, displayed across half of the brain. Finally (from 0:14), the video travels axially through the brain several times, with a color-coding where each color represents the specific view data captured at a specific angle rotation of the sample (18 views in total). The inlet shows the various modalities: arrowheads pointing inward show the lightsheet origin, arrowheads pointing outward show the camera detection position. All arrowheads in red: the information shown combines data from all views. colorscale: a doubled "BIOP12colors" look-up table from Fiji.

### Supplementary Video 2

**Scattered signal by mvLSSM, coronal travel through the mouse brain.** Similar to Supplementary Video 1, the video starts with the autofluorescence signal to position the brain in coronal orientation. At 0:06 the fractional anisotropic is shown through a half brain travel. From 0:10, the video travels coronally through the brain several times, with a color-coding where each color represents the specific view data captured at a specific angle rotation of the sample (18 views in total). The inlet shows the various modalities: arrowheads pointing inward show the lightsheet origin, arrowheads pointing outward show the camera detection position. All arrowheads in red: the information shown combines data from all views. Colorscale: a doubled "BIOP12colors" look-up table from Fiji.

### Supplementary Video 3

**Scattered signal by mvLSSM, sagittal travel through the mouse brain.** Similar to Supplementary Video 1, the video starts with the autofluorescence signal to position the brain in coronal orientation. At 0:11 the fractional anisotropic is shown through a third of brain travel. From 0:14, the video travels coronally through the brain several times, with a color-coding where each color represents the specific view data captured at a specific angle rotation of the sample (18 views in total). The inlet shows the various modalities: arrowheads pointing inward show the lightsheet origin, arrowheads pointing outward show the camera detection position. All arrowheads in red: the information shown combines data from all views. Colorscale: a doubled "BIOP12colors" look-up table from Fiji.

#### Supplementary Video 4

**3D animation illustrating the infinite-cylinder scattering model in two light-sheet microscopy configurations.** Light scattering from cylindrical fibers produces detectable peaks depending on the microscope geometry. (1) macroSPIM configuration: Left-side illumination (yellow arrow, $\mathbf{r}_i$) and back detection (blue arrow and square, $\mathbf{r}_o$). Light is scattered in a cone centered on the fiber axis (orange cone), and a peak is observed when scattered light reaches the detector (bottom-left plot). (2) Ultramiscroscope configuration: Left-side illumination and top detection. The scattering cone remains centered on the fiber axis, but the position, number (one or two) and spacing of the peaks vary with the fiber tilt (α).

#### Supplementary Video 5

**Animation of the triangulation approach for 3D orientation reconstruction.** Example of orientation reconstruction in a case without (1) and with (2) tilt uncertainty. (2). In both cases, in the first mounting position there are 2 fibers that have the same β but different α. After mounting rotation, in case (1) the two fibers have a different β', which enables 3D reconstruction through triangulation. In the case (2), they have the same β' after mounting rotation. Thus, their tilting is not solvable through triangulation.

#### Supplementary Video 6

**Comparison of estimated directions with tLSSM and dMRI.** Axial slices of the estimated directions through RGB-coding with tLSSM (left) and diffusion MRI (right).

#### Supplementary Video 7

**Tract density imaging of the tLSSM mouse brain.** Axial slices of the mapping of 30 million tracts onto the imaging grid generated from the tLSSM directions in the mouse brain. RGB-color coding represents the orientation of the tracts.
